# ZNFX1 uses two-component ubiquitin circuitry to quarantine viral RNA

**DOI:** 10.1101/2025.09.05.674436

**Authors:** Daniel R. Squair, Eilidh Rivers, Hanna Sowar, Arda Balci, Roosa Harmo, David J. Wright, Gaurav Beniwal, Mathieu Soetens, Sunil Mathur, Aidan Tollervey, Callum Stanton, Adam J. Fletcher, Satpal Virdee

## Abstract

The detection of viral RNA inside cells triggers a diverse range of antiviral responses, including global translation inhibition, interferon secretion and RNA sequestration. Mutations in the gene *ZNFX1* cause severe paediatric immunodeficiencies, including chronic viral infection and autoinflammation. Here, we show that ZNFX1 is an RNA helicase with cryptic and unusual bifurcating E3 ubiquitin ligase activity. Nucleotide-dependent RNA binding stimulates ZNFX1 to generate complex ubiquitin chains via a two-component ubiquitin circuit wired in parallel, with ubiquitin flux occurring via either of two competing paths. One route produces K63-linked polyubiquitin that drives ZNFX1 aggregation and RNA entrapment; the other route produces K48-linked polyubiquitin that drives ZNFX1 turnover. RNA entrapment restricts RNA virus replication, and is reversible by deubiquitination. Patient ZNFX1 variants are defective for viral restriction, linking RNA entrapment to antiviral immunity *in vivo*.

## Introduction

Eukaryotic cells encode diverse protein-based mechanisms for detecting virus replication. In the cytosol, retinoic acid inducible gene I (RIG-I)-like receptors (RLR) – members of the RNA helicase super family 2 (SF2) – form filaments on viral dsRNA, inducing the helical tetramerisation of their N-terminal tandem caspase activation and recruitment domains (2CARD). This structural conformation is imprinted on the CARDs of mitochondrial membrane-anchored mitochondrial antiviral signaling protein (MAVS) ^1^, triggering prion-like aggregation of MAVS into a signalosome ^2^. The MAVS platform recruits the kinase TBK1, phosphorylating IRF3 and licencing interferon (IFN) gene transcription ^3^. In these and orthogonal nucleic acid detection systems, molecular clustering is a unifying property that integrates virus detection with signal amplification.

K63-linked polyubiquitin chains (K63-Ub) act as tethers during the assembly of innate immune signaling platforms ^4^. K63-Ub, generated by RLR-associated E3 ubiquitin ligases (E3), including Riplet, TRIM25 and TRIM65, stimulate RNA sensing by promoting RLR 2CARD clustering ^5^. These ‘allosteric’ E3s bind to the RLR helicase core primarily through their PRYSPRY and RING domains ^6,7^, and stimulate the direct transfer of ubiquitin from the loaded E2 enzyme conjugate (E2∼Ub) to the RLR substrate. Of note, the other major class of ubiquitin ligase, the ‘transthiolating’ E3, undergoes thioester exchange with E2∼Ub to form a covalent intermediate. A feature of the “helicase-plus-E3” system is that its function necessitates synchronous expression of two polypeptides, which can be regulated by IFN signaling ^8^. Mutations in genes involved in sensing viral nucleic acids – including RLRs – can predispose individuals to infectious disease, often by otherwise benign pathogens as in the case of Mendelian Susceptibility to Mycobacterial Disease (MSMD) ^9^. These ‘inborn errors of immunity’ (IEI) can also cause broader immunopathologies including chronic viral infections and neuroinflammatory conditions ^10,11^.

Mutations in a poorly understood yet distantly related SF1 RNA helicase, zinc finger NFX1-type containing 1 (*ZNFX1*), across 32 patients from at least 17 unrelated families, have been reported to cause a range of immunopathologies and acute susceptibility to bacterial and viral infections – including MSMD and COVID-19 ^12–17^. Relatedly, ZNFX1 was identified in a screen for genes with antiviral behaviour toward SARS-CoV-2 ^18^. Although not yet well studied, ZNFX1 has been proposed to act as an RNA sensor functionally reminiscent of RLRs ^19^. *ZNFX1* is conserved from invertebrates to mammals ^20^, and in the latter is a ‘core-ISG’ ^8^ (likely reflecting a central role in innate immunity). ZNFX1 is also implicated in stress granule formation and function ^13,21^, mRNA stability ^14,22^, the lysosomal damage response ^21^ and inflammasome activation ^23^; all pathways that intersect with RNA virus replication.

Here, we show that human ZNFX1 is an unannotated and unprecedented bimodal E3, combining the two archetypal E3 mechanisms – allosteric and transthiolating – into a single polypeptide. Bimodal E3 activity is RNA-responsive, bifurcates from a common E2 binding site, is interdependent with the helicase ATPase activity and comprises competing routes for ubiquitin flux leading to the synthesis of distinct polyUb topologies. K63-Ub promotes ZNFX1-mediated aggregation on – and entrapment of – viral RNA, silencing mRNA translation. Thus, ZNFX1 represents the first description of an autonomous immune factor combining helicase and E3 activities to sense and inhibit RNA viruses.

## Results

### ZNFX1 is an interferon-stimulated E3 ligase

We initially carried out activity-based protein profiling (ABPP) to obtain insights into the E3 ligases involved in the IFN response. We used E2∼Ub activity-based probes (ABPs) that covalently trap proteins that undergo thioester exchange with the E2∼Ub conjugate – a hallmark of a transthiolating E3 (**Fig. 1A**) ^24,25^. The workflow entails: preparation of cell extracts; ABP incubation in extracts; enrichment of ABP-labelled proteins; proteomic analysis (**Fig. 1B**). To quantitate changes in protein abundance, we combined ABPP with Stable Isotope Labelling by Amino acids in Cell culture (SILAC-ABPP) ^26^. Upon type I IFN stimulation, both heavy- and light-isotope labelled cells displayed equivalent upregulation of established ISGs, confirming no adverse impact from metabolic labelling on the IFN response (**Supplementary Fig. 1A**). We employed ABPs derived from UBE2D3, one of the four closely related and functionally promiscuous E2 ubiquitin-conjugating enzymes in the UBE2D subfamily (UBE2D1–4) ^27^, and UBE2L3, an E2 that only functions with transthiolating E3s ^28^. and treated heavy and light THP-1 derived macrophages with or without IFN, respectively, for 18 h before harvesting.

**Figure 1.**
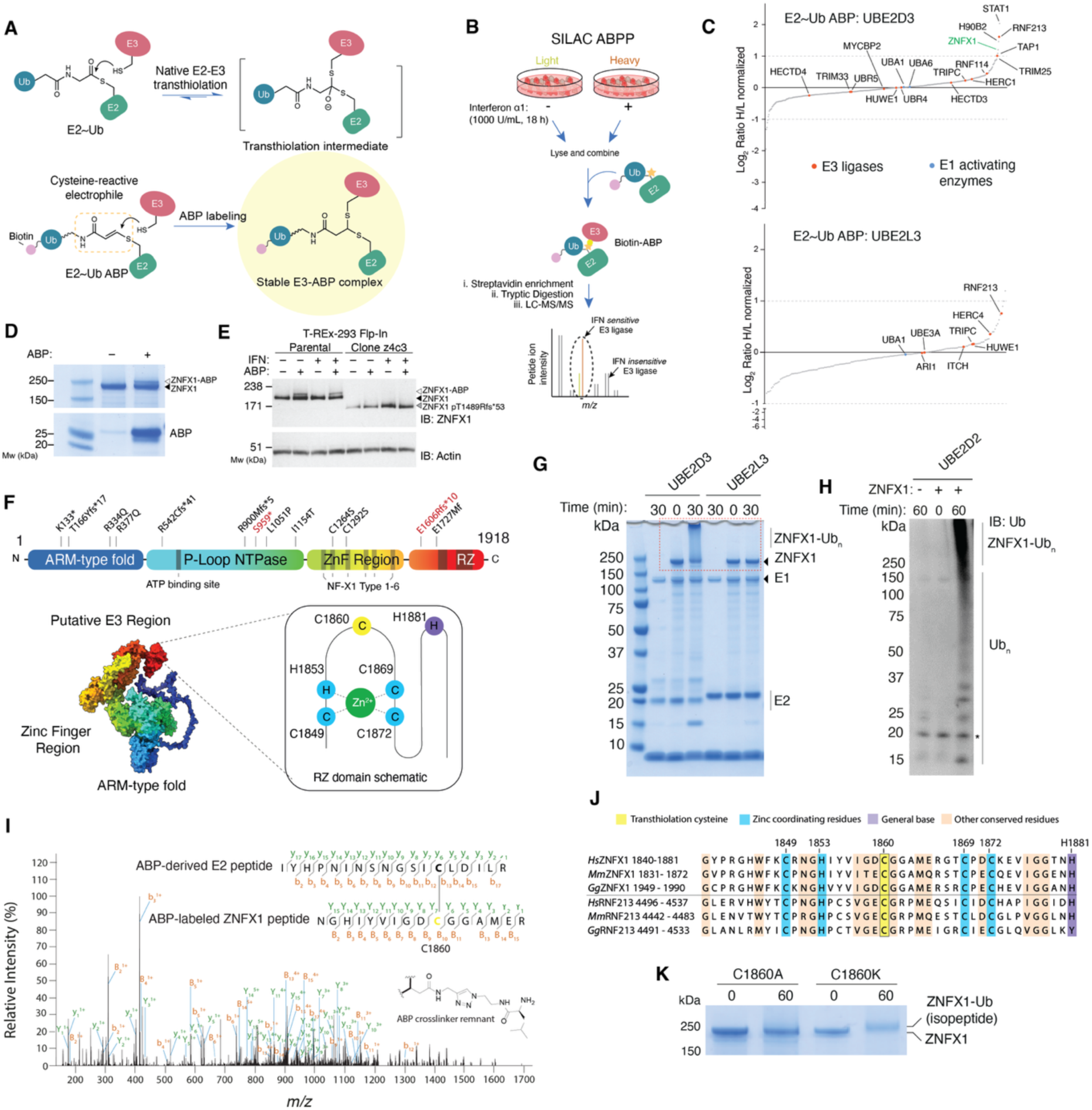
Activity-based profiling identifies ZNFX1 as a transthiolating E3 ligase induced by interferon. **A**) E2∼Ub ABPs form a covalent complex with E3s that covalently receive Ub onto a catalytic cysteine nucleophile through a process termed transthiolation. **B**) Combining ABP profiling with stable isotopic labelling of amino acids in cell culture (SILAC-ABPP) allows proteome-wide quantitation of changes in transthiolating E3 activity in THP-1 cells upon interferon stimulation. **C**) **Top,** waterfall plot depicting the 501 quantitated SILAC-ABPP ratios with the UBE2D3 ABP, which identifies a small cohort of E3 ligases putatively activated by interferon. The RNA helicase ZNFX1 is poorly characterized but has an RZ domain with potential E3 activity. Bottom, **t**he 191 quantitated SILAC-ABPP ratios with the UBE2L3 ABP**. D)** ZNFX1_FL_ (3 µM) purified from insect cells undergoes ABP (25 µM) labelling. **E)** Endogenous ZNFX1 in lysate from T-Rex-293 Flp-in cells in labelled by ABP. A clone lacking the C-terminal 429 residues (z4c3) is not. **F**) ZNFX1 domain architecture and patient variants associated with immune disease. Residue Cys1860 resides within the ZNFX1 RZ domain and has putative catalytic activity. Structure corresponds to Alphafold model. **G**) ZNFX1_FL_ forms high molecular weight species upon addition of ubiquitin cascade components when partnered with UBE2D3 but is inactive with UBE2L3. **H**) Immunoblot analysis demonstrating activity with UBE2D2. Free Ub chains are formed as well as chains that migrate with a molecular weight consistent with ZNFX1_FL_ autoubiquitination. **I)** Representative XL-MS MS^2^ spectrum for a crosslinked peptide corresponding to ABP modification of Cys1860 in ZNFX1. Spectrum corresponds to a 5^+^ precursor ion (theoretical [M+H^+^] = 4038.985; experimental [M+H^+^] = 4038.991). **J**) Residue Cys1860 is in the RZ domain and conserved across ZNFX1 and RNF213 orthologues. His1881 is also conserved with RNF213 where it is a general base essential for E3 activity. Numbering corresponds to human ZNFX1. **K**) Consistent with Cys1860 undergoing transthiolation with E2∼Ub, a ZNFX1_FL_ C1860K mutant forms a stable isopeptide adduct when incubated with ubiquitin cascade components.

For the UBE2D3-derived ABP, 6 proteins were recovered with a >2-fold increase in SILAC-ABPP signal upon IFN treatment (STAT1, RNF213, H90B2, ZNFX1, TAP1 and TRIM25) along with 6 proteins with a >2-fold decrease (**Fig. 1C & Supplementary Table 1**). The E1 activating enzymes UBA1 and UBA6, which are constitutively expressed and active, served as internal controls and, as expected, showed negligible changes in signal. Changes in the recovery of the transthiolating E3s HECTD3, UBR5, MYCBP2, HUWE1, HECTD3, TRIPC and HERC1 were also below our assigned ± 2-fold threshold. The UBE2L3-derived probe showed a distinct profile and no E1 or E3 enzymes were detected in ABP-omitted controls, confirming probe specificity **(Fig. 1C**, **Supplementary Fig. 1B & Supplementary Table 1**).

Further supporting the utility of the SILAC-ABPP methodology, RNF213 is a recently defined transthiolating E3 and an ISG that is functional with UBE2D3 and UBE2L3 ^29,30^. While also an ISG E3, TRIM25 operates via an allosteric mechanism, implying the ABP has covalently attached to TRIM25 at an ‘opportunistic’ cysteine in the vicinity of the TRIM25 RING domain (itself a cysteine-rich fold). In fact, we often detect TRIM25 with UBE2D3- or UBE2L3-based ABPs, suggesting it is a common off-target E3 ^25^. STAT1, an established ISG, was the most highly enriched protein upon IFN stimulation and its specific enrichment with the UBE2D3 probe suggests it can bind this E2. Interestingly, the poorly characterized SF1 RNA helicase ZNFX1 showed an increased SILAC-ABPP signal (2.3-fold) with the UBE2D3 probe, suggesting it might have E3 activity (**Fig. 1C**). Supporting its identification by SILAC-ABPP, both full length ZNFX1 expressed in insect cells (ZNFX1_FL_) and cellular ZNFX1 underwent labelling with the ABP (**Fig. 1D,E**). Serendipitously, during efforts to derive a *ZNFX1* deleted T-Rex-293 Flp-In cell line using CRISPR/Cas9, we generated a homozygous clone expressing ZNFX1 lacking the C-terminal 429 residues (**Supplementary Fig. 1C**). Unlike the full-length protein, truncated ZNFX1 was not labelled by ABP (**Fig. 1E**), indicating a determinant for ABP reactivity lies within this region.

ZNFX1 is a poorly understood enzyme but is a conserved ISG ^8^ and has been proposed to function as a sensor of viral RNA ^19^. Moreover, patients harbouring non-synonymous mutations in *ZNFX1* suffer a range of severe immunopathologies (**Fig. 1F**) ^12–14^. Although ZNFX1 is not known to function as an E3, it shares an RNF213-ZNFX1 (RZ) domain with RNF213, where it is required for thioester-mediated ubiquitin transfer (**Fig. 1F & Supplementary Fig. 1D**) ^29,31^. However, the E2 binding site in RNF213 comprises a C-terminal domain (CTD) that is absent from ZNFX1 ^29^, raising questions about how ZNFX1 might recruit an E2∼Ub conjugate. The RZ domain resides very near to the ZNFX1 C-terminus, potentially explaining the lack of labelling in the truncated cellular mutant (**Fig. 1G,H**).

Many E3s undergo auto-modification when incubated with components of the ubiquitin cascade, which can be used to validate E3 activity. To assess E3 ligase activity, ZNFX1_FL_ was tested with the E2 enzymes UBE2D3 and UBE2L3. ZNFX1_FL_ generated free ubiquitin chains and a high-molecular-weight smear, suggestive of polyubiquitination in an E1-dependent manner, but only in the presence of UBE2D3 **(Fig. 1I, J and Supplementary Fig. 1E)**. Coincident depletion of the ZNFX1 band is consistent with substantial auto-modification with polyubiquitin chains. We next tested a panel of 29 E2s and observed activity with several E2s, which was most robust with UBE2D1-4, which can function with both allosteric and transthiolating E3s, including RNF213 (**Supplementary Fig. 1F-I)** ^27,32^. Inactivity with UBE2L3 was in contrast to RNF213, in alignment with the SILAC-ABPP specificity profile **(Fig. 1C, D)**, and supports the notion that ZNFX1 has unique E2 and catalytic requirements.

We sought to identify the cysteine labelled by the ABP – that should correspond to an essential active site residue – allowing both E3 mechanism and E3-dependent functions to be investigated. Proteolytic digestion of the ABP-adduct, observed for ZNFX1_FL_, yields a cross-linked species containing the corresponding E3 peptide, which can be identified by crosslinking mass spectrometry (XL-MS) (**Supplementary Fig. 1J**) ^24,25,29,33^. XL-MS spectra converged on a peptide labeled at Cys1860, located within the RZ domain, which is analogous to the active site cysteine in RNF213 and is conserved across ZNFX1 orthologues (**Fig. 1K,L & Supplementary Table 2**). In further support of C1860 undergoing transthiolation with E2∼Ub, a ZNFX1_FL_ C1860K – but not a C1860A – mutant trapped the otherwise transient E3∼Ub catalytic intermediate, presumably as a stable isopeptide conjugate (**Fig. 1M**) ^34^. Collectively, these data reveal that ZNFX1 is an E3 ligase that utilises a catalytic cysteine in its RZ domain to accept ubiquitin from the E2∼Ub conjugate.

### ZNFX1 restricts viral replication in an E3-dependent manner

ZNFX1 is reported to bind incoming viral RNA and induce IFN secretion to reduce viral replication ^19^. To understand whether RZ-dependent E3 activity of ZNFX1 contributes to this function, we first sought an assay for ZNFX1 antiviral activity. In a recent screen for cellular proteins recruited to RNA viral genomes, we noticed that ZNFX1 was amongst a series of proteins recruited to Sindbis virus (SINV), a positive-sense ssRNA virus from the *Togaviridae* family ^35,36^ (**Supplementary Fig. 2A**) SINV replicates efficiently in HEK293 cells, so we generated a complete *ZNFX1* knock-out-reconstitution system in T-REx-293 Flp-In cells, to study the effects of novel ZNFX1 variants on viral replication (**Supplementary Fig. 2B**). We employed an established reverse genetics system to generate fluorescent protein-encoding recombinant SINV RNA genomes by *in vitro* transcription ^35^. High-titre SINV was then harvested from host cells following viral RNA (vRNA) transfection. To distinguish antiviral phenotypes acting early or late during vRNA replication, we used two recombinant viruses encoding fluorescent proteins expressed at either early (mScarlet, fused to nSP3) or late (mCherry, under the control of a subgenomic promoter) stages of viral replication (**Fig. 2A**) ^35^. These viruses allow longitudinal, non-invasive fluorescence detection of SINV replication, and facilitate the measurement of viral replication kinetics in the presence and absence of ZNFX1. Following infection of T-REx-293^ZNFX1^ ^KO^ cells, mScarlet signals were detected first, followed by mCherry signals ∼5 h later, consistent with early and late gene expression, respectively (**Fig. 2B,C**). Markedly, expression of ZNFX1 caused a ∼5 h delay in mScarlet and mCherry detection (**Fig. 2B,C**). To test whether this restriction was via an induction of IFN secretion, as suggested ^19^, we also treated cells with ruxolitinib, a JAK inhibitor that disables IFN signaling through the JAK-STAT pathway. However, ruxolitinib had minimal effect on SINV replication kinetics, regardless of ZNFX1 expression (**Supplementary Fig. 2C,D**). This suggested that ZNFX1 can also exhibit direct antiviral activity, which occurs soon after viral entry.

**Figure 2.**
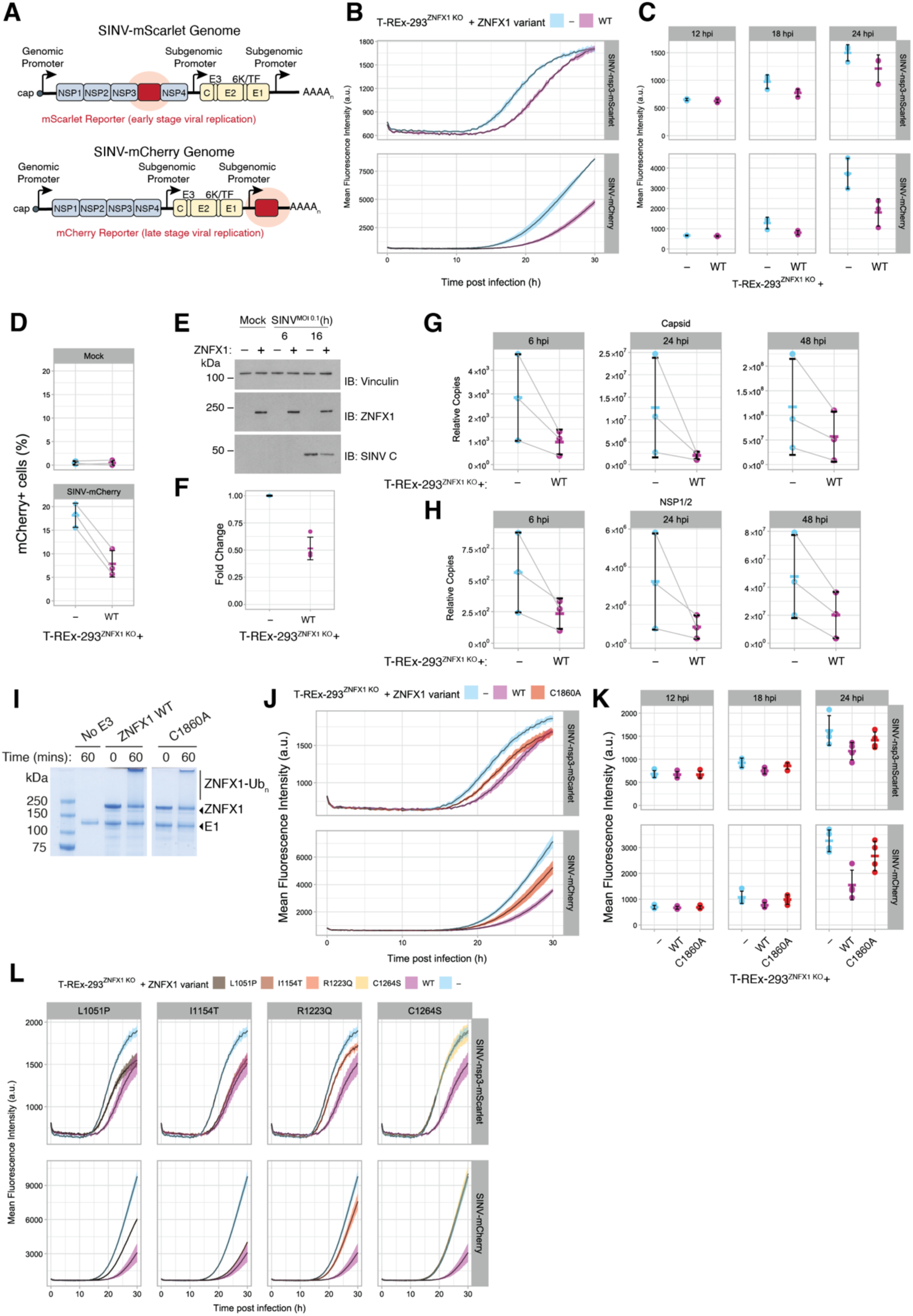
Optimal ZNFX1 viral restriction requires transthiolating E3 activity. **A**) Two SINV reporter viruses were used in this study. The first expresses mScarlet tagged to NSP3, under the control of the genomic promoter. The second expresses mCherry from a second subgenomic promoter. **B**) Red fluorescent signal in T-REx-293^ZNFX1^ ^KO^ cells (blue) reconstituted with WT-ZNFX1 (pink) and infected with SINV-nsp3-mScarlet (top panel) or SINV-mCherry (bottom panel) (MOI = 0.1). Fluorescence was measured at intervals of 15 min in an incubated plate. Data are presented as mean values ± SD from 3 independent infections. **C**) Mean ± SD of red fluorescent signal from three biological replicates of (B) at 12, 18, and 24 hours post infection. **D**) Percent of mCherry+ cells (T-REx-293^ZNFX1^ ^KO^ (blue) reconstituted with WT-ZNFX1 (pink)) following mock infection or infection with SINV-mCherry (MOI = 3.75x10^-3^) for 24 h. **E**) SINV Capsid protein levels were analyzed in T-REx-293^ZNFX1^ ^KO^ cells and T-REx-293^ZNFX1^ ^KO^ cells reconstituted with WT-ZNFX1 following infection with WT SINV (MOI = 0.1) for 0, 6, and 16 hours. Vinculin served as a loading control. **F**) Mean ± SD of the level of SINV Capsid protein quantified from Western blots (E) and normalised to the level in the KO sample of each repeat (n = 3). **G, H**) Quantification of SINV capsid (G) and NSP1/2 (H) transcripts in T-REx-293^ZNFX1^ ^KO^ cells (blue) reconstituted with WT-ZNFX1 (pink) infected with WT SINV (MOI = 0.01) for 6, 24, or 48 h by RT-qPCR. **I**) Mutation of Cys1860 in ZNFX1FL reduces, rather than ablates, autoubiquitination. **J**) Red fluorescent signal in T-REx-293^ZNFX1^ ^KO^ cells (blue) reconstituted with WT-ZNFX1 (pink) or C1860A-ZNFX1 (red) infected with SINV-nsp3-mScarlet (top panel) or SINV-mCherry (bottom panel) (MOI = 0.1). Fluorescence was measured at intervals of 15 min in an incubated plate reader (37°C and 5% CO2). Data are presented as mean values ± SD from 3 independent infections. **K**) Mean ± SD of red fluorescent signal from **four** biological replicates of (J) at 12, 18, and 24 hours post infection. **L**) Red fluorescent signal in T-REx-293^ZNFX1^ ^KO^ cells (blue) reconstituted with WT-ZNFX1 (pink) or a ZNFX1 mutation identified in a patient, infected with SINV-nsp3-mScarlet (top panel) or SINV-mCherry (bottom panel) (MOI = 0.1). Fluorescence was measured at intervals of 15 min in an incubated plate reader (37°C and 5% CO2). Data are presented as mean values ± SD from 3 independent infections.

The antiviral phenotype was quantified by flow cytometry (**Fig. 2D**), and was not exclusive to mutant viral genomes as wild type SINV was also restricted by ZNFX1; Western blots (**Fig. 2E,F**) and RT-qPCR (**Fig. 2G,H**) for viral capsid protein (SINV C) or genomes, respectively, revealed commensurate reductions in replication. Reductions in genomic or genomic + subgenomic vRNA were observed from as early as 6 h post infection (hpi), consistent with an IFN-independent restriction mechanism (**Fig. 2G,H**). ZNFX1 expression had no effect on cell doubling rate, indicating that differences in viral growth were not attributable to differences in cell density (**Supplementary Fig. 2E**). We also found that the negative-sense RNA virus human metapneumovirus, encoding GFP (hMPV-GFP), replicated with delayed kinetics in ZNFX1-expressing cells (**Supplementary Fig. 2F**), suggesting broad viral targeting by ZNFX1 ^19^. Combined, these data support a model whereby ZNFX1 inhibits viral replication through an unknown pathway that operates in parallel to its described effects on IFN signalling.

We next assessed the requirement for transthiolating E3 activity in ZNFX1 antiviral behaviour. To test that C1860A substitution abolishes ZNFX1 E3 activity, we measured E3 activity of a ZNFX1_FL_ C1860A mutant. Unexpectedly, E3 activity was reduced rather than ablated (**Fig. 2I**), suggesting additional determinants of ZNFX1 E3 activity exist. This stands in contrast to RNF213, where equivalent mutation of the RZ abolishes E3 activity ^29^. We then expressed the C1860A variant in T-REx-293^ZNFX1^ ^KO^ cells, infecting cells with SINV variants. While the WT protein delayed SINV replication by ∼ 5 h as before, ZNFX1 C1860A restriction was less potent, delaying replication by ∼ 2.5 h – manifest as an intermediate phenotype between the knockout and ZNFX1 WT-expressing cells (**Fig. 2J,K**). Combined, these data demonstrate that ZNFX1 is a direct-acting antiviral protein whose restriction potency tracks with E3 activity. Moreover, while transthiolating activity plays a contributory role, an additional property of ZNFX1 is necessary for optimal defence.

Finally, we sought to test whether this antiviral function correlated with the discordant immune response in children bearing *ZNFX1* mutations. We selected 4 non-synonymous mutations (i.e. that did not cause protein truncation) in or near the helicase domain – L1051P, I1154T, R1223Q, C1264S ^12,14^ – and expressed these in the T-REx-293^ZNFX1^ ^KO^ background, confirming expression by Western blot (**Supplementary Fig. 2G**); use of a KO background allowed us to mimic the effect of patient homozygosity. We then compared SINV replication in these and the parental KO and ZNFX1 WT-expressing cells. Whereas I1154T had little effect on ZNFX1 control of viral replication, substitutions L1051P and R1223Q caused substantial loss of restriction activity, while C1264S was completely inactive (**Fig. 2L**). Whilst both L1051P and R1223Q were homozygous in patients ^12,14^, I1154T occurred in heterozygotes alongside C1292S; our data suggest the latter might in fact be the deleterious mutation ^14^. C1264S also occurred in two heterozygotes with truncated variant E1727Kfs*11 ^14^; in this case, the latter mutant might not be pathogenic in isolation. Strikingly, the antiviral phenotypes in our assay correlated with clinical outcomes. Patients carrying L1051P and R1223Q substitutions, while experiencing severe neurological symptoms and blood/liver disorders, were still alive at 33 months and 6 years of age, respectively, when reported ^12,14^. In contrast, and supporting the importance of residue C1264, both children carrying the C1264S change suffered hemophagocytic lymphohistiocystosis-like disease characterised by multi-organ failures, RNA and DNA viral infections, and passed away at 3 and 16 months of age ^14^. Thus, these data suggest that the cellular antiviral assay reveals correlates of ZNFX1 immune function *in vivo*, encouraging us to pursue its enzymatic mechanism in detail.

### ZNFX1 exhibits defining features of both transthiolating and allosteric E3s to generate complex ubiquitin chain architecture

Invariably, the other known transthiolating E3s – HECTs, RBRs, RCRs and RNF213 – are wholly dependent on their catalytic cysteines for E3 activity ^25,28,29,37^. In contrast, ZNFX1 is an RZ-type E3 that retains partial activity when the catalytic cysteine is substituted. To understand the biochemical basis for this unusual behaviour, we sought to dissect the complete mechanism of ZNFX1-mediated ubiquitin transfer. Allosteric E3s catalyze Ub transfer from a more reactive ‘closed’ E2∼Ub conformation whereas the transthiolating HECT and RBR E3s do not ^28,38–42^. E2 residues N77, D87, and L104 (UBE2D3 numbering) are essential for allosteric activation, as they promote and stabilize the closed E2∼Ub conformation; in contrast, N114 is partially dispensable ^28,43–47,38,39,48,49^ (**Fig. 3A & Supplementary Fig. 3A**). E3 activity was abolished when E2 variants containing substitutions at essential residues were combined with ZNFX1, whilst the N114Q variant supported an intermediate level of activity (**Fig. 3B & Supplementary Fig. 3B**). Collectively, these data demonstrate that ZNFX1 E3 activity is strictly dependent on the closed E2∼Ub conformation. The only example of a transthiolating E3 sharing this requirement is the RCR E3 MYCBP2 ^34^.

**Figure 3.**
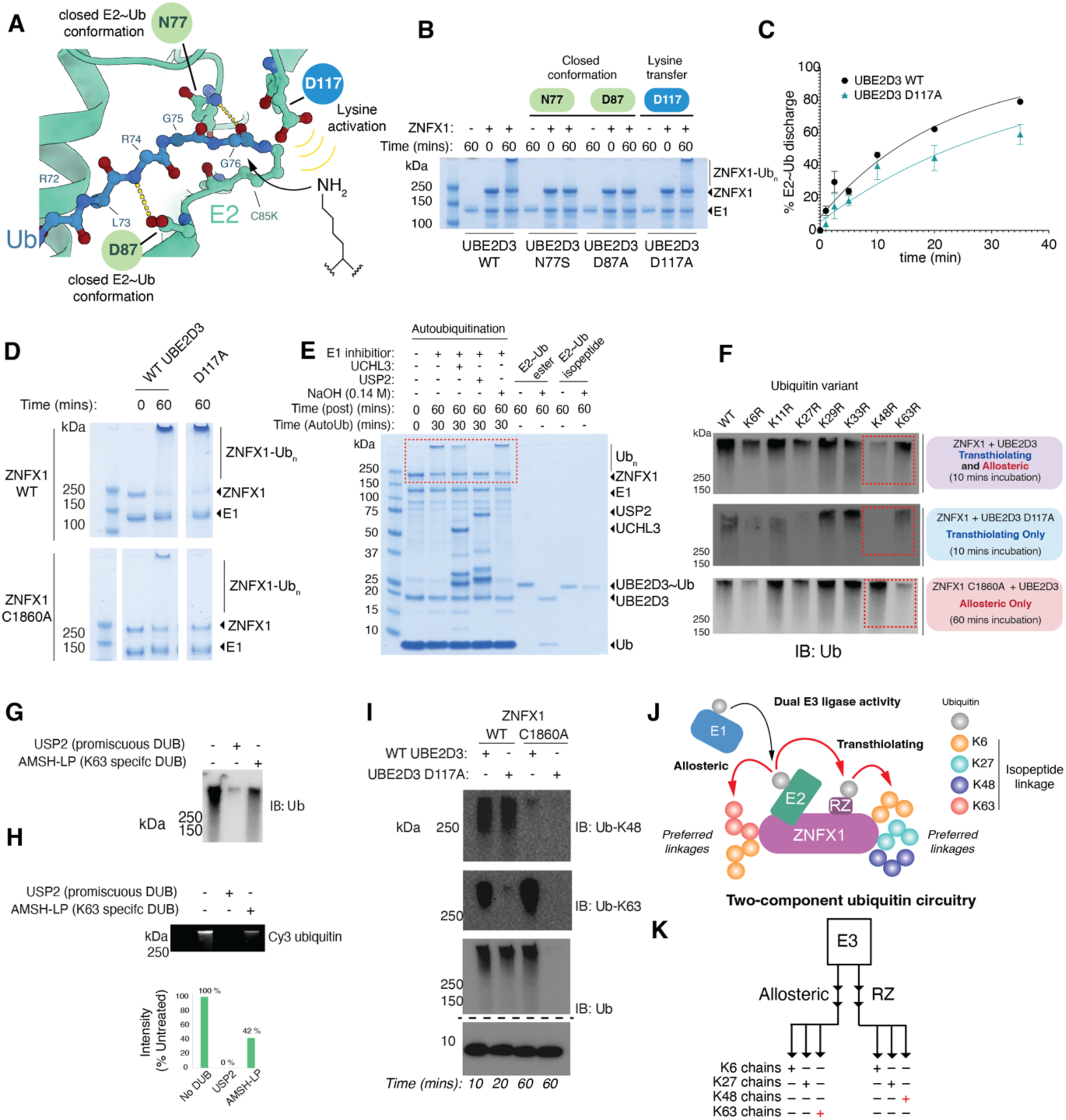
ZNFX1 assembles distinct ubiquitin chains with dual E3 activity. **A**) E3s that require a closed E2∼Ub conformation are not functional with N77 or D87 mutations. Allosteric E3s that catalyze aminolysis are not functional when residue D117 is mutated. Coordinates taken from PDB ID 4AP4 ^38^. **B**) ZNFX1 is inactive with the N77S and D87A variants of UBE2D3 but retains activity with D117A, consistent with it undergoing transthiolation with a closed E2∼Ub conformation. **C**) Single turnover E2∼Ub discharge assay for ZNFX1_FL_ combined with WT and D117A variants of UBE2D3. E2 variants were preloaded with Ub, followed by E1 inhibition ^25^. The kinetics of ZNFX1_FL_ E3 activity were determined by fitting the discharge of Ub from E2∼Ub as a function of time to a single exponential function (*n* = 2). **D**) Activity of ZNFX1 WT and C1860A when partnered with WT and D117A variants of UBE2D3. Extended panel shown in Supplementary Fig. 3B. **E**) High molecular weight ZNFX1_FL_ species are sensitive to the promiscuous DUB USP2, but not NaOH, indicative of them containing amide-linked ubiquitination. The high molecular weight species is partially sensitive to UCH-L3 treatment. **F**) Assessment of ubiquitin linkage specificity assembled by ZNFX1_FL_ (top panel) reveals that formation of high molecular weight ubiquitin species is most sensitive to K6R, K27R, and K48R substitutions. A similar pattern is observed when ZNFX1 is restricted to transthiolating activity (middle panel). In contrast, when ZNFX1 is limited to allosteric activity, the formation of high molecular weight ubiquitin species is most sensitive to K6R, K27R, and K63R substitutions (bottom panel). **G**) Ubiquitination reactions with wild type components were treated with the promiscuous DUB USP2 or the K63-specific DUB AMSH-LP. Reactions were visualized by anti-Ub immunoblot. **H**) Reactions were also performed with Cy3-labelled Ub and visualized by in-gel fluorescence. **I)** Distinct assembly of Lys48 and Lys63 linkages by transthiolating and allosteric activities, respectively, was confirmed with native ubiquitin and linkage-specific antibodies. **J**) Schematic depicting the dual E3 mechanism and the preferred ubiquitin linkages each activity assembles. **K)** The dual E3 activity is akin to parallel wiring in an electrical circuit.

Another E2 mutant we used for diagnostic purposes is D117A, which ablates transfer to lysine but not cysteine residues (**Fig. 3A**). In accordance, transthiolating E3s – including MYCBP2 – retain full activity with this variant (**Supplementary Fig. 3A**) ^34,40^. Unexpectedly, instead of a benign effect, we observed a 2-fold reduction in ZNFX1 reaction kinetics, further implying that ZNFX1 has both allosteric and transthiolating E3 activities (**Fig. 3C**). Confirming the presence of allosteric activity, the signal was lost when we combined ZNFX1_FL_ C1860A with UBE2D3 D117A (**Fig. 3D**). Furthermore, in alignment with the RZ-dependent transthiolating activity sharing RNF213’s requirement for a general histidine base, activity was abolished when ZNFX1_FL_ H1881N was partnered with UBE2D3 D117A (**Fig. 1K & Supplementary Fig. 3C**). These data demonstrate that ZNFX1 is an E3 ligase that utilizes independent transthiolating and allosteric activities – an unprecedented property for an E3.

RZ-dependent lipid A ubiquitination by RNF213 might occur on hydroxyl residues, as it can be hydrolysed under high pH conditions ^31^. However, ZNFX1-dependent high-molecular weight polyubiquitin smears – indicative of autoubiquitination – were resistant to sodium hydroxide treatment but could be depleted by the deubiquitinase (DUB) USP2, indicating amide-linked, rather than ester-linked, ubiquitin chains predominate in this protein-only assay (**Fig. 3E**). The DUB UCHL3, which cannot cleave ubiquitin homopolymers, preferring disordered sites, partially removed the polyubiquitinated species, supporting a degree of autoubiquitination at intrinsically disordered regions (IDRs) of ZNFX1 ^50^ (**Fig. 3E**).

We reasoned that allosteric and transthiolating activities might perform distinct biochemical functions. The ubiquitin system employs diverse ubiquitin chain architectures to encode defined cellular signals ^51^. Using a panel of ubiquitin variants each lacking one of the seven lysine residues required for canonical chain synthesis (K6R, K11R, K27R, K29R, K33R, K48R and K63R), we assessed the overall linkage profile of ZNFX1. Bulk ZNFX1_FL_ autoubiquitination was most impaired with K48R, K27R and K6R mutants, suggesting these linkages are predominant (**Fig. 3F upper panel**). By coupling ZNFX1 WT with UBE2D3 D117A, and ZNFX1 C1860A with UBE2D3 WT, isolated transthiolating and allosteric mechanisms, respectively, could also be profiled. We observed that both activities were sensitive to K6R and K27R Ub, implying these atypical linkages are a product of both transthiolating and allosteric activities (**Fig. 3F middle and lower panels**). However, only the transthiolating activity was impaired with the Ub K48R mutation (**Fig. 3F, middle panel**), whilst isolated allosteric activity was uniquely impaired with K63R Ub (**Fig. 3F bottom panel**). To assess K63 chain abundance under native reaction conditions, we treated samples with the K63-specific DUB AMSH-LP ^52^. This reduced the high-molecular-weight ubiquitin signal by ∼50% (**Fig. 3G**), and a similar depletion was observed by in-gel fluorescence with labeled Ub (**Fig. 3H**), demonstrating that K63 linkages are a substantial product of normal ZNFX1 E3 activity. We confirmed the distinct preferences for K48 and K63 linkages were maintained with wild-type Ub using linkage-specific antibodies (**Fig. 3I**). Thus, the two ZNFX1 E3 ligase mechanisms independently catalyse the synthesis of alternative polyubiquitin chains; the allosteric mechanism generates K63-linked chains, while the RZ generates K48-linked chains (**Fig. 3J**). Interestingly, K48-linked chains are typically associated with degradative outcomes, while K63-linked chains are prominent within immune signalling pathways in a non-degradative role ^5,53^. This ability to operate as a two-component ubiquitin circuit might grant ZNFX1 with functional diversity (**Fig. 3K**).

### ZNFX1 bimodal E3 activity emerges from a novel RING-like domain

We next asked whether ZNFX1’s independent E3 activities emanate from distinct E3 modules or bifurcate from a common E2 docking site. Unlike RZ E3 counterpart RNF213, ZNFX1 possesses neither RING nor CTD and has no other annotated E2 binding domain. To identify the E2 binding site/s in ZNFX1, we generated an *in silico* ZNFX1-UBE2D2 complex with the AlphaFold2 Multimer protein fold tool ^54^. UBE2D2 was predicted to interact with high confidence at an unannotated region of ZNFX1, with the interaction centered around E2 residue F62 contacting ZNFX1 residue F1575. (**Fig. 4A and Supplementary Fig. 4A**) ^34,41,55,56^. Alanine substitution of either residue in ZNFX1_FL_ or UBE2D2 abolished E3 activity, suggesting these two residues might experience π-stacking interactions and revealing that the observed dual E3 activity bifurcated from a single E2 binding site (**Fig 4B & Supplementary Fig. 4B**). Interestingly, a glutathione S-transferase (GST)-tagged construct encompassing this region of ZNFX1, but lacking the RZ domain (residues 1462-1652, ZNFX1_E2BR_), was more active than ZNFX1_FL_ C1860A, indicating allosteric activity is suppressed in the context of the full-length protein (**Fig 4B,C**). In further support of ZNFX1 possessing allosteric activity, ZNFX1_E2BR_ was labelled by a photo-reactive ABP – that assesses an E3’s ability to stabilize the closed E2∼Ub conformation – in an F62-dependent manner (**Supplementary Fig. 4C**) ^57^. Consistent with the full-length protein, ubiquitination activity of ZNFX1_E2BR_ was also abolished by F1575A substitution (**Supplementary Fig. 4D**).

**Figure 4.**
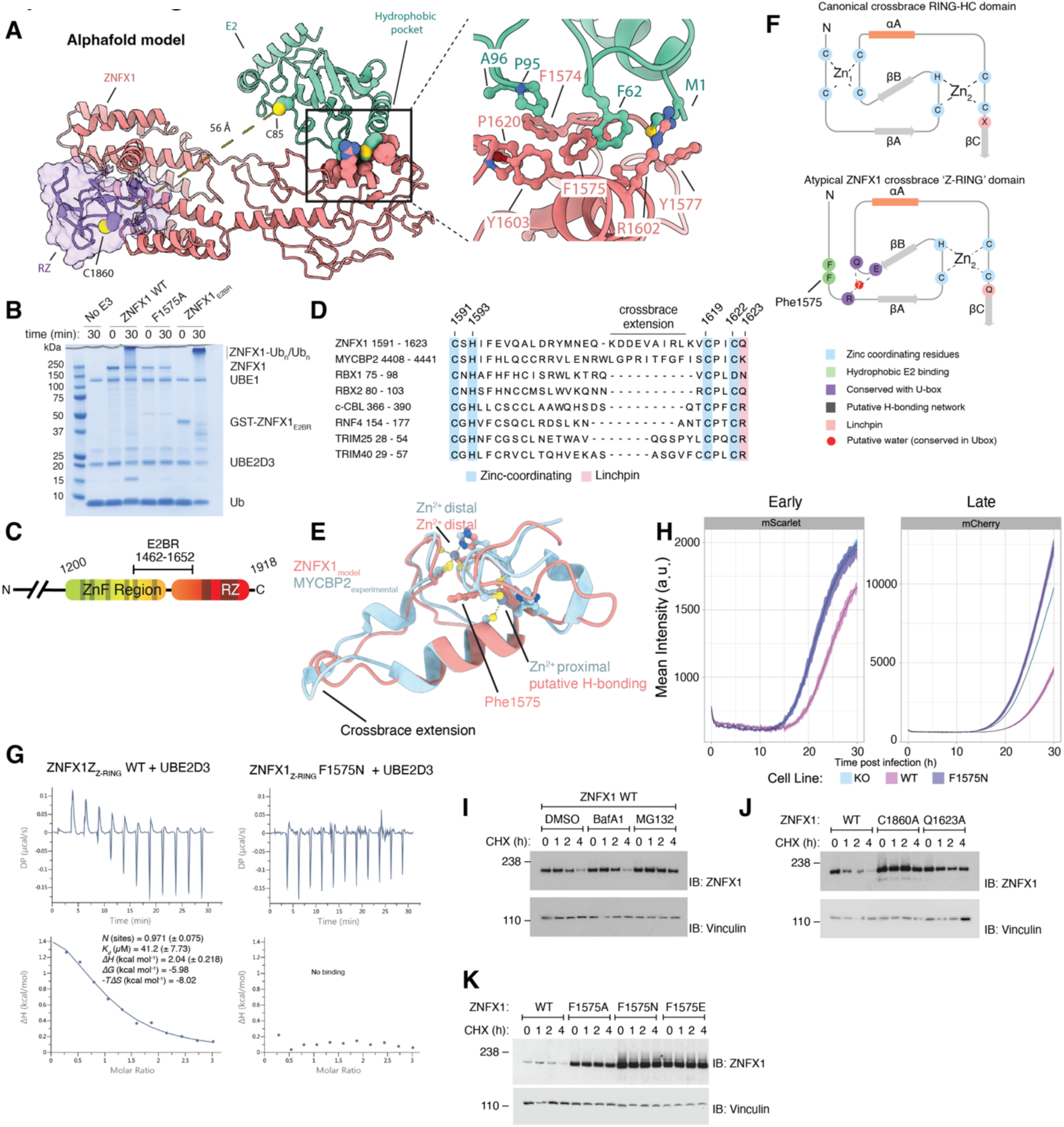
Dual E3 activity bifurcates from an atypical E2 binding region and is essential for viral restriction. **A**) Model of a ZNFX1-UBE2D3 complex calculated with Alphafold2 Multimer. Inset is a close up of the predicted interface between E2 and ZNFX1 residue F1575. **B**) ZNFX1_FL_ autoubiquitination is undetectable when F1575 is mutated, showing that dual E3 activity bifurcates from a common E2 binding module. ZNFX1 lacking the RZ domain (GST-ZNFX1_E2BR_) undergoes robust autoubiquitination. **C**) Residue boundaries for the E2BR (E2 binding region) construct that lacks the RZ domain. **D**) Sequence alignment of the E2 binding ‘Z-RING’ with canonical crossbrace RING domains. It shares a loop extension found in the MYCBP2 RING domain. **E**) Structural alignment of the Alphafold model of the Z-RING with the crystal structure of the MYCBP2 RING domain (PDB ID: 5O6C). **F**) Schematic representation of the canonical RING-HC and Z-RING domain. **G**) Isothermal titration calorimetry was used to measure the affinity between the predicted Z-RING and UBE2D3. Binding is entropically driven and undetectable with a F1575N mutant. Errors are from data fitting from a single experiment, but a parallel experiment yielded similar values. **H)** Red fluorescent signal in T-REx-293^ZNFX1^ ^KO^ cells (blue) reconstituted with WT-ZNFX1 (pink) or F1575A-ZNFX1 (purple) infected with SINV-nsp3-mScarlet (top panel) or SINV-mCherry (bottom panel) (MOI = 0.1). Fluorescence was measured at intervals of 15 min in an incubated plate reader (37°C and 5% CO_2_). **I**) Western blot analysis of ZNFX1 in T-REx-293^ZNFX1^ ^KO^ reconstituted with WT-ZNFX1 and treated with DMSO, BafA1 (100 nM), or MG132 (10 µM) and cycloheximide (0.1 mg/mL) for 0, 1, 2, or 4 h. Vinculin served as a loading control**. J,K)** Western blot analysis of ZNFX1 in T-REx-293^ZNFX1^ ^KO^ cells reconstituted with WT, C1860A, Q1623A, F1575A, F1575N or F1575E ZNFX1 and treated with cycloheximide (0.1 mg/ml) for 0, 1, 2, or 4 h. Vinculin served as a loading control.

The most prevalent E2-binding motif in the human genome is the Really Interesting New Gene (RING) domain, characterized by a cross-brace architecture that coordinates two zinc ions, positioned either proximally or distally relative to the bound E2 residue F62 ^58^. Sequence alignments of the region encompassing the essential F1575 residue with diverse RING domains highlighted four conserved ZNFX1 residues (C1591, H1593, C1619 and C1622) that could serve as a Zn^2+^ coordination site (**Fig. 4D**). Structural alignment with the MYCBP2 RING domain suggested that the ZNFX1 E2 binding module may also adopt cross-brace architecture, where the distal Zn^2+^ coordinating site is present, but the proximal site is absent, seemingly replaced by a hydrogen bonding network (**Fig. 4D-F, Supplementary Figure 4E-G**). This architecture is reminiscent of the SP-RING domain found in SUMO E3 ligases and the inverse configuration found in the UBR4 hemiRING domain ^59,60^. We purified a small construct corresponding to this putative cross-brace region - we termed the ‘Z-RING’ (residues 1533-1631, ZNFX1_Z-RING_) – and measured E2 binding by isothermal titration calorimetry (ITC). UBE2D3 underwent entropically driven binding (*K_d_* = 41 (±8) µM; -*TΔS* = -8.02 kcal mol^-1^; Δ*H* = +2.04 kcal mol^-1^) – consistent with the affinity range for established E3s (**Fig. 4G**) ^58^. No binding was detected with an F1575N mutation, providing further credence to the *in silico* predicted E2 binding site.

We next tested the model that ZNFX1 cellular activity tracks with E3 activity by comparing SINV replication kinetics in cells lacking or expressing WT or F1575N ZNFX1. As before, WT ZNFX1 expression delayed SINV replication by ∼ 5 h. Strikingly, ZNFX1 F1575N failed to restrict viral replication (**Fig. 4H, Supplementary Fig. 4H**), suggesting dual ubiquitination activity, bifurcating from the Z-RING, underlies the observed antiviral phenotype.

Typically, a linchpin residue is needed for allosteric RING E3 activity, which locks the E2∼Ub into the reactive closed conformation ^39^. While a basic arginine residue is optimal and found in ∼50% of RING domains, suboptimal glutamine, asparagine or lysine residues are also utilized ^61^. Sequence alignments highlighted Q1623 as the analogous residue in ZNFX1 (**Fig. 4D,F**). Consistent with it having a functional role, substitution of Q1623 with alanine led to reduced activity with ZNFX1_FL_, with a more pronounced reduction observed for the allosteric-only ZNFX1_E2BR_ construct – suggestive of allosteric activity being more dependent on the functional integrity of the linchpin residue than transthiolating activity (**Supplementary Fig. 4I, D**). Furthermore, *in silico* modelling of UBE2L3 with ZNFX1_E2BR_, coupled with E2 mutational analysis, provided insight into their functional incompatibility – UBE2L3 residue Lys96 likely causing a steric clash with the Z-RING (**Supplementary Fig. 3B**) ^34^. Taken together, these data support the conclusion that the Z-RING forms an atypical cross-brace fold with intrinsic allosteric E3 activity. In accordance, the residues proposed to have structural and functional roles are conserved across ZNFX1 orthologues (**Supplementary Fig. 4J**).

As an assay for cellular activity, we reasoned the dual E3 activity that assembles complex polyUb chain architectures contributes to ZNFX1 degradation ^51^. In line with this, WT ZNFX1 had a cellular half-life of ∼ 2 h that was extended by proteasome but not lysosome inhibition (**Fig. 4I**). By using proteasomal degradation as a proxy for E3 activity, we compared the stability of ZNFX1 variants. Supporting its role in generating K48-linked chains, transthiolation mutation C1860A improved stability (**Fig. 4J**). A modest stabilisation by linchpin substitution Q1623A suggests the ZNFX1 allosteric activity might modulate turnover. As expected, F1575X (X = A/N/E) substitutions abolished ZNFX1 turnover (**Fig. 4K**), aligning with a loss of total ubiquitination activity. Together these data reveal that ZNFX1 dual E3 activities emanate from a novel RING-like domain and that RZ-catalysed K48-linked chains drive ZNFX1 turnover.

### RNA binding regulates ZNFX1 E3 ligase activity

Having established that ZNFX1 cellular stability inversely correlates with E3 catalytic integrity, we next asked whether RNA binding affects E3 activity. In general, RNA helicases make contacts with the sugar-phosphate backbones of RNA, affording relative degrees of non-specific RNA binding. To explore this further, we first sought ZNFX1 residues required for RNA binding. The ZNFX1 AlphaFold structure had high confidence metrics in the helicase domain (pLDDT scores > 70) (**Supplementary Fig. 5A**), suggesting it faithfully captured a relevant protein fold. To confirm this, we superposed this structure with a crystal structure of the helicase domain of prototypic SF1 member Upf1 – an RNA helicase involved in mRNA nonsense-mediated decay (NMD) ^62,63^ – in complex with single-stranded polyU and ADP:AlF_4_^−^ (a mimic of an ATP hydrolysis intermediate) ^64^. The two helicase domains aligned well (root mean square deviation (RMSD) value between 246 pruned atom pairs = 2.77 Å) (**Supplementary Fig. 5B**). Using this structural alignment, we identified a series of ZNFX1 residues that were potential sites of RNA contact (**Supplementary Table 3**). Importantly, several of these residues have been characterised for RNA binding in Upf1.

To identify a ZNFX1 mutant defective in RNA binding we generated a panel of ZNFX1 variants mutated throughout the putative RNA binding site. We then developed a cellular RNA binding assay using biotin-labelled poly(I:C), a dsRNA mimic that serves as a functional ligand for both SF1 and SF2 helicases, including RIG-I and ZNFX1 ^19,65^. Biotin-poly(I:C) was incubated in cell extracts from T-REx-293^ZNFX1^ ^KO^ cells expressing inducible ZNFX1 variants. WT ZNFX1 was enriched by biotin-poly(I:C), confirming ZNFX1 binds directly to RNA (**Supplementary Fig. 5C**). Importantly, a variant harbouring double mutations at positions F1210 and R1216 (F1210A/R1216A), conserved across ZNFX1 and Upf1 orthologues and referred to as the FARA mutant, was not enriched by biotin-poly(I:C), suggesting a specific defect in RNA binding and that ZNFX1 binds RNA via its conserved SF1 helicase domain (**Fig. 5A,B & Supplementary Fig. 5C**). Interestingly, the FARA variant was highly stabilised in cells, even after extended periods of translation inhibition (**Fig. 5C**), supporting a model of ZNFX1 cellular stability regulated by endogenous RNA-dependent E3 activity.

**Figure 5.**
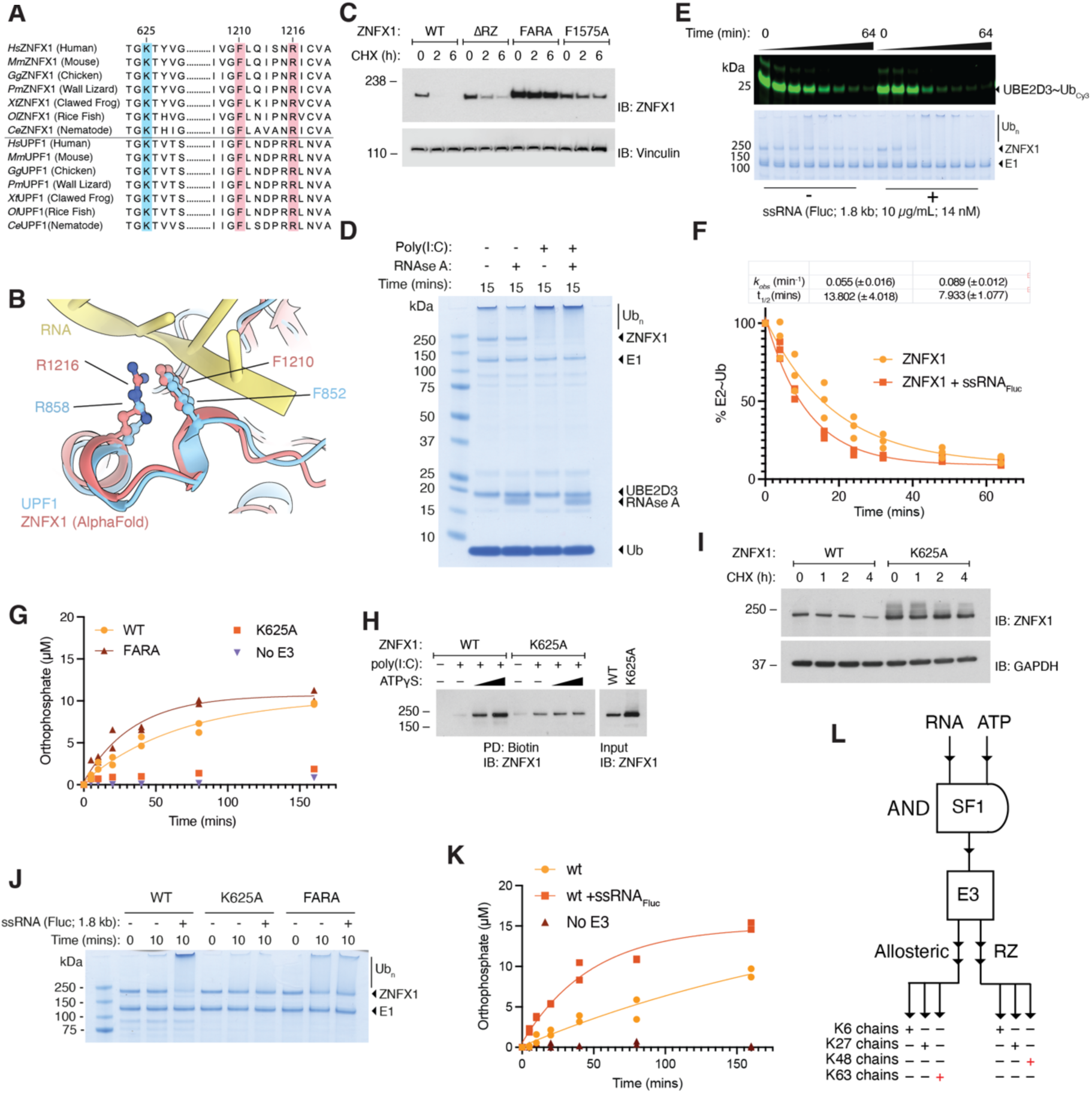
ZNFX1 demonstrates nucleotide-dependent RNA binding that enhances NTPase and E3 activity. **A)** Conservation of residues shown to be required for RNA binding between ZNFX1 and Upf1. Sequences were aligned with the Clustal algorithm. **B**) Superposition of the ZNFX1 AlphaFold model with the crystal structure of Upf1 bound to a ATP hydrolysis intermediate mimic (PDB ID: 2XZO) ^64^. **C**) Western blot analysis of ZNFX1 in T-REx-293^ZNFX1^ ^KO^ cells reconstituted with WT, ΔRZ, FARA, or F1575A-ZNFX1 and treated with cycloheximide (0.1 mg/mL) for 0, 2, or 6 h. **D)** ZNFX1_FL_ E3 assay in the absence and presence of poly(I:C). RNase A cleaves at cytosine bases, generating ssRNA. **E**) Single-turnover E2∼Ub discharge assay measuring E3 activity in the presence and absence of ssRNA_Fluc._ **F**) Quantitation of E2∼Ub discharge shown in F). In-gel fluorescence was quantified and fitted to a single exponential decay function (*n* = 3). **G)** Malachite green ATPase assay for ZNFX1_FL_ defective in RNA (FARA) and nucleotide (K625A) binding. Orphophosphate production was fitted to a single exponential decay function (*n* = 2). **H)** Enrichment of ZNFX1 by biotin-poly(I:C) in the presence of increasing concentrations of ATPgS. Nucleotide dependent binding to poly(I:C) is lost upon mutation of the Walker A P-loop residue K625. **I**) Western blot analysis of ZNFX1 in T-REx-293^ZNFX1^ ^KO^ cells reconstituted with WT or K625A-ZNFX1 and treated with cycloheximide (0.1 mg/mL) for 0, 1, 2, or 4 h. GAPDH served as a loading control. Mutation of the Walker A P-loop residue K625 stabilizes cellular ZNFX1. **J**) E3 activity assay for different ZNFX1 variants in the presence and absence of an in vitro transcribed 1.8 kb ssRNA molecule (ssRNA_Fluc_). Mutation of the Walker A P-loop residue K625 ablates RNA responsiveness. The RNA-binding defective ZNFX1 FARA variant is also unresponsive to ssRNA_Fluc_ but has elevated E3 activity. **K**) Malachite green ATPase assay for ZNFX1 in the presence and absence of ssRNA_Fluc_ (*n* = 2). **L)** The requirement for two distinct signals to regulate bifurcated E3 ligase activity resembles an AND logic gate, coordinating a parallel ubiquitin signaling circuit with both shared and unique outputs.

We next investigated the effect of RNA binding on ZNFX1 E3 activity biochemically. In accordance with dsRNA stimulating E3 activity, poly(I:C) potently enhanced the accumulation of high molecular weight species, and coincident ZNFX1 band depletion (**Fig. 5D**). Only high molecular weight forms of poly(I:C) enhanced ZNFX1 depletion (**Supplementary Fig. 5D**), consistent with a reported preference to bind long versus short dsRNA ^19^. Conversely, dsDNA-mimic poly(dA:dT), uncut plasmid DNA or short ssDNA oligonucleotides had no effect, indicating activation was RNA-specific (**Supplementary Fig. 5D,F**). Interestingly, the poly(I:C)-enhanced ubiquitination remained following pretreatment of poly(I:C) with RNase A (**Fig. 5D**), which selectively cleaves 3’ of pyrimidine nucleotides, suggesting ssRNA (polyI) is sufficient to activate the ZNFX1 E3 ^66^. To confirm ssRNA was sufficient, we obtained defined ssRNA molecules by *in vitro* transcription, generating 1.8 kb ssRNAs derived from either an arbitrary DNA template (ssRNA_Fluc_) or 1.8 and 4.0 kb ssRNAs derived from *PRKAA2* cDNA (ssRNA*_PRKAA2_*_-S_ and ssRNA*_PRKAA2_*_-L_) (**Supplementary Fig. 5E**). ZNFX1 was shown to bind the latter to support autophagy and anti-*Mycobacterium tuberculosis* immunity ^22^. Addition of either ssRNA_Fluc_ or ssRNA*_PRKAA2_*_-L_ also potentiated formation of high molecular weight ZNFX1 species with coincident depletion of free ZNFX1 (**Supplementary Fig. 5G,H**), confirming ZNFX1 displays no sequence specificity under these conditions. A modest “hook” effect was apparent with an ssRNA*_PRKAA2_*_-L_ dose-response, suggesting avidity-driven binding of ZNFX1 to RNA enhances its activity (**Supplementary Fig. 5G**). To quantify E3 stimulation by ssRNA binding, we used single-turnover E2∼Ub discharge assays, recording a two-fold increase in ubiquitin discharge by ZNFX1 in the presence of ssRNA_Fluc_ (**Fig 5E,F**). Combined, these data reveal that ZNFX1 is an ssRNA-activated E3.

### ZNFX1 ATP hydrolysis and RNA binding are interconnected

ATP binding to SF1 and SF2 family helicases can bring both RecA domains into a closed conformation to reveal an RNA-binding cleft ^64^. As P-loop NTPases, these enzymes contain a highly conserved Walker A motif that regulates ATP binding by coordinating the γ-phosphate. Returning to the ZNFX1-Upf1 structural alignment, ZNFX1 residue K625 superposed precisely with Upf1 K498, identifying the Walker A motif in ZNFX1 (**Supplementary Fig. 5I & Table 3**). Confirming this, *in vitro* ATP consumption by ZNFX1_FL_ was abolished in a K625A mutant (**Fig. 5G**). To test whether ATP binding influences ZNFX1 RNA binding, we first measured binding between biotin-poly(I:C) and ZNFX1 in cell extracts, in the presence or absence of poorly hydrolyzable ATP analogue ATPγS. Indeed, ATPγS strongly enhanced WT ZNFX1 recovery with biotin-poly(I:C), but not for the Walker A mutant K625A (**Fig. 5H**), confirming ATPγS operated via the predicted position. In support of E3 activity being dependent on ATP-dependent RNA binding, the cellular stability of ZNFX1 K625A was also substantially enhanced compared to WT protein (**Fig. 5I**), indicating a directionality from ATP binding to RNA engagement to E3 activity.

In agreement with this model, neither ZNFX1_FL_ K625A nor FARA mutants were responsive to ssRNA in *in vitro* ubiquitination assays (**Fig. 5J**). However, conversion of ZNFX1_FL_ FARA to high molecular weight species was deregulated, occurring even in the absence of ssRNA (**Fig 5J**). Interestingly, the FARA variant also demonstrated elevated basal ATPase activity (**Fig 5G**), suggesting a complex cross-talk between RNA binding and ATPase activity. ZNFX1 ATPase activity was also robustly stimulated by the presence of ssRNA_Fluc._ (**Fig. 5K**), suggesting that ZNFX1 undergoes RNA translocation and RNA-binding augments both of ZNFX1’s catalytic activities. Combined, these data suggest that ATP hydrolysis and RNA binding by ZNFX1 are highly cooperative processes, as seen in other RNA helicases ^63^. Sequential rounds of ATP binding and hydrolysis by ZNFX1 may promote its clustering or translocation along RNA, thereby activating its E3 ligase function. Thus, the requirement for both RNA and ATP acts as an upstream ‘AND gate’ that potentiates the two-component ubiquitin circuit driven by ZNFX1’s bifurcated E3 activity (**Fig. 5L**).

### Ubiquitin-regulated RNA entrapment by ZNFX1 aggregates

Our data so far suggest that ZNFX1 binds ATP and RNA to activate its E3-dependent restriction of viral replication. To test this model in cells, we assessed SINV replication kinetics with ZNFX1 variants deficient in ATP binding (K625A) or RNA binding (FARA). As before, expression of wild-type ZNFX1 reduced the replication kinetics of both fluorescent SINV mutants. In contrast, virus restriction was completely abrogated in cells expressing the K625A or FARA mutants **(Fig. 6A,B; Supplementary Fig. 6A)**, indicating that the helicase domain functions upstream of E3 activity. These results support a model in which ATP-dependent RNA binding is essential for ZNFX1 engagement with viral RNA and antiviral function in cells.

**Figure 6.**
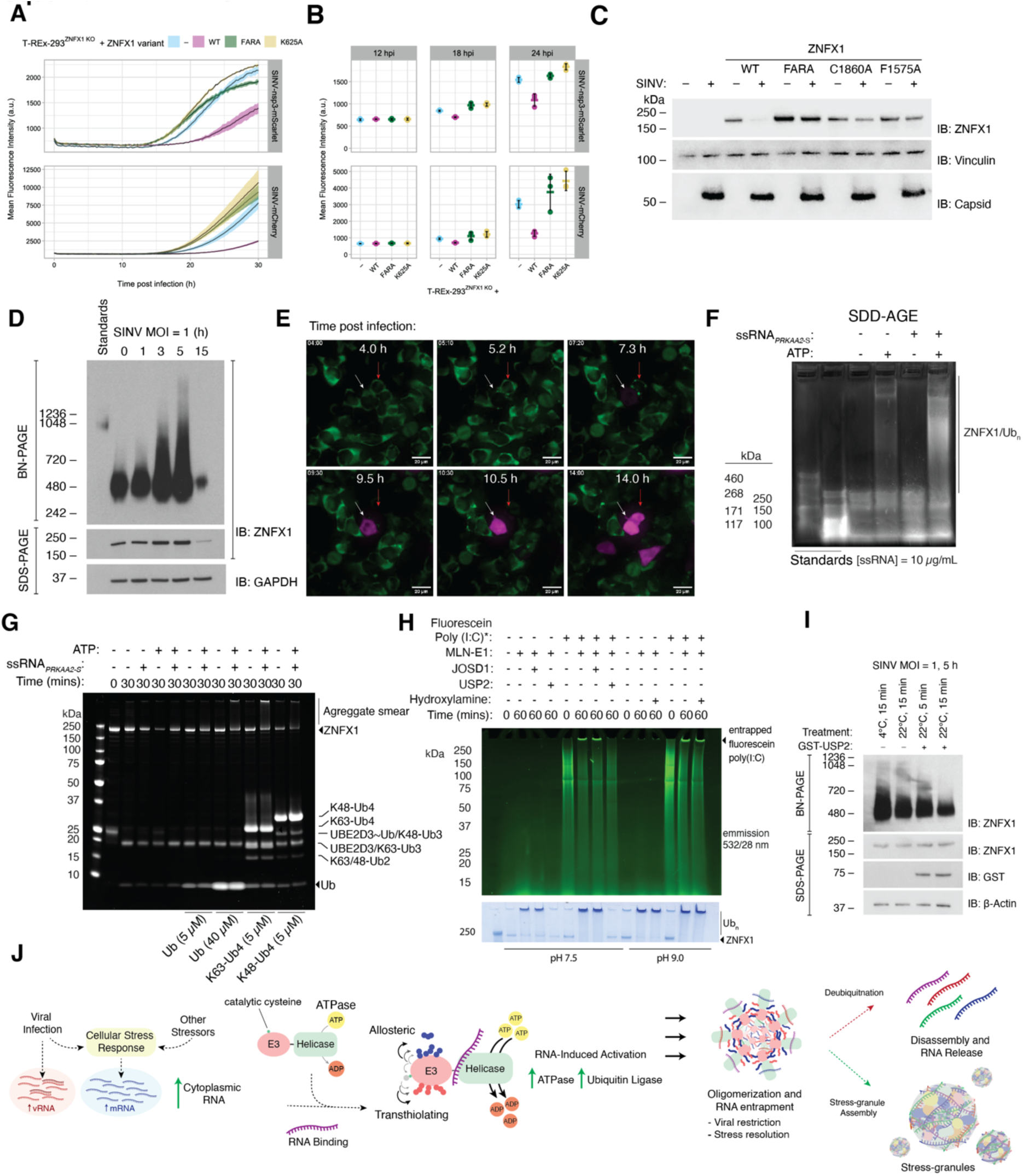
RNA binding and nucleotide binding are both essential for viral restriction which is mediated by RNA entrapment. Red fluorescent signal in T-REx-293^ZNFX1^ ^KO^ cells (blue) reconstituted with WT (pink), FARA (green), or K625A (yellow) ZNFX1 and infected with SINV-nsp3-mScarlet (top panel) or SINV-mCherry (bottom panel) (MOI = 0.1). Fluorescence was measured at intervals of 15 min in an incubated plate reader (37°C and 5% CO_2_). Data are presented as mean values ± SD from 3 independent infections. **B)** Mean ± SD of red fluorescent signal from three biological replicates of (A) at 12, 18, and 24 hours post infection. **C)** Western blot analysis of ZNFX1 in T-REx-293^ZNFX1^ ^KO^ cells reconstituted with WT, FARA, C1860A, or F1575A ZNFX1 and mock infected or infected with WT-SINV for 18 h (MOI = 1). Vinculin served as a loading control. **D)** Blue Native PAGE gel (top panel) and SDS-PAGE gel (bottom panels) of in T-REx-293^ZNFX1^ ^KO^ cells reconstituted with WT-ZNFX1 and infected with WT SINV (MOI = 1) for 0, 1, 3, 5, or 15 h. **E)** Stills from live cell confocal microscopy analysis of T-REx-293^ZNFX1^ ^KO^ cells reconstituted with mNeonGreen-ZNFX1 WT, infected with SINV-mCherry (MOI = 20). Times indicate hours post infection. White and red arrows are fixed on two individual cells. **F)** SDD-AGE of ZNFX1 autoubiquitination reactions in the presence and absence of ATP and ssRNA. Reactions were run for 30 min at 37 °C, resolved by Coomassie-stained agarose gel, and imaged by fluorescence (Cy5 emission filter). High molecular weight, SDS-resistant ZNFX1 species were markedly enhanced by ssRNA. Aggregates migrated above the 460 kDa ladder limit, with apparent molecular weights exceeding 1 MDa. **G)** ZNFX1_FL_ aggregation is triggered by a 5-fold molar excess of preloaded E2∼Ub, revealing that the high molecular weight ZNFX1 species induced by ssRNA (generated by treatment of poly(I:C) with RNase A), consists of autoubiquitinated and unmodified forms of ZNFX1. Additionally, unanchored ubiquitin enhances the formation of high molecular aggregates, with K63-linked tetraubiquitin exerting a stronger effect than either K48-linked tetraubiquitin or monoubiquitin. Aggregation is driven by cooperative inputs from RNA, ATP, E3 ligase activity, and unanchored ubiquitin. Shown is a Coomassie-stained SDS–PAGE gel imaged using far-red detection. **H)** Fluorescein-labelled poly(I:C) accumulates in the well during ZNFX1 autoubiquitination, indicating co-aggregation of RNA and protein. ZNFX1 aggregates are dissolved by the broad-specificity DUB USP2, but not by the DUB JOSD1 or hydroxylamine, which cleave ester linkages. Gel was imaged by in-gel fluorescence (top) or Coomassie staining (bottom). **I)** Blue Native PAGE gel (top panel) and SDS-PAGE gel (bottom panels) of in T-REx-293^ZNFX1^ ^KO^ cells reconstituted with WT-ZNFX1 and infected with WT SINV (MOI = 1) for 5 h. The lysates were incubated at 22°C (room temperature) with or without the deubiquitinase USP2 for 5 or 15 min. **J)** Model of RNA- and ubiquitin-induced ZNFX1 aggregation. Cellular stimuli that increase RNA levels, such as viral infection, promote RNA binding by ZNFX1 via its helicase domain in an ATP-dependent manner. RNA engagement enhances the ubiquitination and ATPase activities of ZNFX1, driving its oligomerisation and entrapment of RNA. These aggregates may act as precursors to stress granules or be disassembled by deubiquitinases.

To understand the restriction mechanism further, we next turned our attention to ZNFX1 itself, asking how the protein responds to viral infection. After 18 h infection, SINV infection induced robust depletion of ZNFX1 WT, but not of ZNFX1 variants FARA, C1860A or F1575A (**Fig. 6C**), suggesting that infection had activated the E3 leading to proteasome-dependent degradation. However, under these conditions, we also observed potent and global SINV-induced translational shut-off, as reported ^67^ (**Supplementary Fig. S6B**). Experiments to rescue ZNFX1 expression during infection – using proteasome inhibitors – were obfuscated by the observation that SINV requires proteasome activity for efficient replication (**Supplementary Fig. 6C**). Thus, we were unable to exclude the possibility that ZNFX1 degradation after 18 h was due to non-specific translation inhibition. However, blue native-PAGE (BN-PAGE) analyses revealed that, specifically at short intervals post-infection, ZNFX1 exhibited gradual accumulation into high molecular weight species, which were not reflected by altered migration patterns in SDS-PAGE, implying these species represented higher order assemblies rather than modified ZNFX1 species (**Fig. 6D**). Moreover, the fastest migrating band ran slightly lower than the 480 kDa marker, suggesting ZNFX1 forms a homodimer in uninfected cells (**Fig. 6D**). The cellular aggregation of ZNFX1 was mirrored by a subtle stabilisation of ZNFX1 by SDS-PAGE (**Fig. 6D**). We postulate this is triggered by binding viral RNA because ZNFX1 oligomerisation could also be induced by incubation of poly(I:C) in cell extracts (**Supplementary Fig. 6D**). We also infected cells expressing mNeonGreen-tagged ZNFX1 (mNG-ZNFX1) with SINV-mCherry to perform live cell microscopy analyses, observing a transition of mNG-ZNFX1 from diffuse to punctate expression that preceded the appearance of mCherry protein, supportive of infection-induced ZNFX1 aggregation (**Fig. 6E**). In cells where ZNFX1 aggregation was evident, it occurred on a similar timescale to the aggregates observed by native PAGE analysis (**Fig. 6D**), while ZNFX1 disappearance following aggregation mirrored the degradation observed in Western blots (**Fig. 6C,D**). Thus, SINV infection induces first oligomerisation, then degradation (direct or indirect), of cellular ZNFX1.

We next tested if the high molecular weight species formed during our previous *in vitro* ubiquitination reactions, which up to this point we had considered solely products of extensive autoubiquitination, might also contain higher-order ZNFX1 oligomers. To gauge the molecular weight of the these poorly electrophoresing species, we analysed them by semi-denaturing detergent-agarose gel electrophoresis (SDD-AGE) – a technique developed for prion-like aggregates ^68^. Indeed, ZNFX1 resolved with an electrophoretic mobility lower than that imparted by autoubiquitination alone and oligomer formation was stimulated by ssRNA (**Fig. 6F**). Remarkably, these species were detectable even when boiled in the presence of 4% lithium dodecyl sulfate (LDS), reminiscent of prion-like aggregates (**Fig. 6F**) ^2^. This finding suggested that ssRNA drives ZNFX1 oligomerisation in cells and *in vitro*. To formally exclude the possibility that these aggregates were only composed of extensively autoubiquitinated ZNFX1 species, we limited the number of ubiquitin discharge cycles - and thus the extent of ZNFX1 autoubiquitination - by providing purified, preloaded E2∼Ub (5 µM) instead of E1 and excess free ubiquitin. Once again, inordinately high molecular weight species formed but only in the presence of unanchored Ub at concentrations comparable to those in our conventional E3 assays (40 µM) (**Fig. 6G**). The addition of tetrameric K63-linked Ub had a more potent effect than monomeric Ub, whereas a K48-linked tetramer was comparable to monomeric Ub (**Fig. 6G**). A requirement for E3-dependent Ub conjugation was confirmed as aggregation was not observed when E2∼Ub was omitted (**Supplementary Fig. 6E**). Thus, ATP-dependent RNA binding, E3 activity and unanchored K63-Ub are all required for optimal aggregate formation *in vitro*. This also implies that the allosteric component of ZNFX1’s bifurcated E3 activity not only modulates turnover, but also aggregate formation.

ZNFX1 has been proposed to buffer the immune system to ensure a balanced interferon response to RNA viruses^14^. For example, ZNFX1 stabilizes ISG mRNA but the mechanism remains unexplored. We reasoned RNA entrapment within the observed aggregates might underlie how ZNFX1 acts as a molecular buffer, effecting viral restriction and RNA homeostasis more broadly. To test this, we stimulated autoubiquitination assays with fluorescently labelled poly(I:C). Consistent with RNA entrapment, an LDS-resistant fluorescent signal overlapped with the high molecular weight aggregate (**Fig. 6H**). We considered an alternative explanation for the high molecular weight RNA – namely, that it undergoes direct ubiquitination. To test this, we pretreated poly(I:C) with RNase A generating single-stranded poly(I), which would only contain hydroxyl nucleophilic acceptors demanding base-labile ester-linked ubiquitination that is sensitive to hydroxylamine, and potentially the esterase-specific deubiquitinase (DUB) JOSD1 ^69,70^. However, the RNA signal was resistant to either treatment (**Fig 6H**). Further excluding direct RNA ubiquitination, we failed to detect ubiquitination of RNA nucleotides by LC-MS (**Supplementary Fig. 7A-C**). The RNA entrapped aggregate was insensitive to subsequent RNase A treatment, indicating that once formed, either its integrity was RNA-independent, or it sequestered RNA so effectively that RNA was inaccessible to soluble RNase (**Supplementary Fig. 7D**).

Because ubiquitin conjugation was key to aggregate assembly, we tested whether its removal might result in aggregate dissolution and RNA release. Remarkably, the promiscuous DUB USP2 dissolved the high molecular weight LDS-resistant aggregates, which was presumably associated with the release of the entrapped RNA (**Fig. 6H**). Confirming an analogous ubiquitin-dependent aggregation had occurred during viral infection, we also treated cell extracts from SINV-infected ZNFX1-expressing cells with USP2, which caused the collapse of cellular ZNFX1 oligomers (**Fig. 6I**). In contrast to those formed in vitro, these aggregates were inherently unstable as incubation of extracts at 22 °C for 15 min also caused partial collapse. This could be accounted for by the opposing activity of cellular DUB(s) or other factors present in the native cell extract (**Fig. 6I**). Collectively, these data reveal that RNA binding to ZNFX1 stimulates ZNFX1 E3 and NTPase activity, enabling the entrapment of RNA into ubiquitin-switchable oligomers/aggregates (**Fig. 6J**).

## Discussion

Herein, we report the discovery of a new addition to the expanding cohort of atypical E3 ubiquitin ligases. ZNFX1 functions through the cooperative activity of its E3 module and SF1-type helicase domain, a hitherto undescribed pairing in an E3 ligase. Moreover, the E3 module is unprecedented in the sense that it is a true mechanistic hybrid possessing both allosteric (RING-like) and transthiolating (HECT-like) activities. The regulation of E3 ligase activity by an ATP-dependent catalytic module is a shared feature of RZ domain-containing E3s, as RNF213 transthiolation is activated by ATP binding to its AAA ATPase core^29^. This exposes an emerging paradigm where divergent catalytic modules with nucleotide substrates control the activity of atypical E3 ligase modules within the same polypeptide. In THP-1 cells, RNF213 and ZNFX1 generated the highest transthiolating activity-based probe signals upon IFN treatment, suggesting the RZ domain is highly important for pathogen defence. Supporting this, RNF213 and ZNFX1 belong to a small set of ‘core ISGs’, namely the 60+ genes upregulated by IFN signaling from mammals to birds^8^.

However, despite these commonalities, the NTPase-helicase domain in ZNFX1, and bifurcated E3 activity, set it apart from RNF213. Here, the RZ fold represents an alternative route for ubiquitin flux, in a system akin to a two-component electrical circuit with parallel routes through which current can flow. This novel form of ubiquitin circuitry raises the possibility of regulating ubiquitin outputs in a context-dependent manner. Like a parallel electrical circuit, if one branch is broken, the current will continue in the other; removal of the RZ route potentiated flux down the K63-biased allosteric route, while removal of the allosteric linchpin switched the current through the K48-biased RZ. In this way, either the linchpin, or the RZ, may act as molecular switches to adjust current direction. In the absence of ssRNA, ubiquitin current passed predominantly through the RZ domain, supporting a model where ubiquitin flux might be modulated by transitions in protein conformation, as could occur upon engaging RNA. Furthermore, synchronous K48- and K63-linked chains may act in concert to promote the formation of branched K48/K63 chains. Interestingly, K48/K63 chains accumulate upon inhibition of VCP/p97 – a Ub-dependent segregase – that may therefore facilitate the extraction of ZNFX1 from RNA-induced aggregates ^71,72^.

Importantly, this helicase-dependent E3 ligase mounts a direct antiviral response by sequestering and silencing viral RNA via molecular crowding. Our data reveal a complex interplay between ATP-dependent RNA binding to the ZNFX1 helicase and two-component E3 activity. These two enzymatic functions collaborate to carry out polyubiquitin synthesis and entrapment of viral RNA in ZNFX1 oligomers. Strikingly, we find that these oligomers can be dissolved – and the entangled RNA presumably released – by deubiquitination, even from LDS-resistant aggregates formed *in vitro*, suggesting a mechanism of virus restriction via a poorly soluble condensate or ‘web of ubiquitin’. We also find that unanchored K63-linked chains potentiate ZNFX1-RNA oligomer formation. Whilst we cannot exclude the possibility that aggregation might be seeded by distinct K63-Ub-specific E3s – as occurs for canonical RLR signaling^6^ – the simplest model is that, through intrinsic allosteric E3 activity, ZNFX1 can generate the ubiquitin ‘glue’ autonomously. Therefore, we propose that ZNFX1 is added to the cohort of E3s employing K63-Ub to drive molecular clustering in response to viral RNA^5^.

This mechanism appears to be critical for a balanced immune response. ZNFX1 also binds cellular RNAs^22^, and has been proposed to buffer both foreign and host RNA (i.e. ISG mRNA) allowing not only viral defence but also resolution of immune responses ^14^. Mutations in human *ZNFX1* – now documented in 32 children – appear to be highly deleterious, predisposing their carriers to a variety of immune disfunctions including acute susceptibility to viral infections, neurological disorders and a hemophagocytic lymphohistiocystosis-like immunodeficiency associated with multi-organ failure ^12,14^. Of the four *ZNFX1* patient variants tested for viral restriction in this study, three were functionally impaired. Interestingly, an inverse correlation between antiviral potency *in vitro* and pathology *in vivo* suggests that RNA traps represent a hitherto overlooked yet critical aspect of innate immunity. Thus, our data provide a mechanistic explanation for the recently uncovered *ZNFX1*-linked immunodeficiencies and suggest that deeper understanding of ubiquitin-mediated RNA silencing could inform host-directed therapeutic strategies.

We note that ZNFX1 aggregates bear hallmarks of cellular mRNA/protein condensates known as stress granules (SG), which form following detection of viral RNA by protein kinase R (PKR) and subsequent translation inhibition ^73^, and can have direct antiviral roles through the sequestration of viral RNA ^74^. Oligomeric proteins bound to RNA can contribute to the condensation process – a process termed percolation ^75^. Hence, RNA bound ZNFX1 oligomers might seed the assembly of SGs, while the reversibility of aggregate formation by DUB activity may allow SG disassembly. Indeed, SG assembly is ATP-dependent, and disassembly is coordinated by multiple RNA-binding ATPases and E3 ligases, the latter including TRIM25, TRIM56 and DTX3L ^76^, and ZNFX1 is also described to localise to SGs and induce them upon overexpression ^13,21^. Thus, ZNFX1 could be an E3 that promotes SG assembly, rather than disassembly ^77,78^. However, RNA binding is not a necessity for oligomerisation. The RNA-binding-defective FARA mutant, which exhibits hyperactive ATPase and E3 activity, spontaneously forms oligomers *in vitro*. RNase treatment of pre-assembled aggregates did not induce disassembly. This suggests that while RNA stimulates E3 activity and templated assembly, spontaneous oligomerization can occur in deregulated mutants. The lack of antiviral activity and hyperstability exhibited by the FARA mutant in cells indicates that aggregates on their own are insufficient for restriction, further supporting a model of viral RNA entrapment as the determinant of antiviral behaviour and ZNFX1 turnover. Although a direct role for ATP hydrolysis activity in oligomerization has not been definitively established, its stimulation by RNA binding implies it is a necessary step - potentially for ATP-dependent RNA unwinding or translocation.

In summary, this study identifies ZNFX1 as a novel atypical E3 ubiquitin ligase that uniquely integrates an SF1-type helicase domain with bifurcated E3 activity, functioning through both allosteric and transthiolating mechanisms. Unlike canonical E3s, ZNFX1’s activity is modulated by ATP-dependent RNA binding, enabling it to reversibly entrap viral RNA within oligomers that may be associated with stress granule formation. These aggregates, formed through a ‘ubiquitin web’, represent a distinct antiviral mechanism that does not rely on secondary immune signaling. The discovery that ZNFX1 mutations impair viral restriction provides a mechanistic explanation for *ZNFX1*-linked immunodeficiency and reveals a broader principle: that ubiquitin-mediated RNA entrapment may serve as a direct antiviral strategy, offering potential for therapeutic exploitation.

## Acknowledgments

We thank the MRC Protein phosphorylation and ubiquitylation unit mass spectrometry team and the CVR-Bioimaging Facility, for their support and assistance in this work. S.V., D.R.S, D.J.W., M.S., S.M. and C.S. are funded by funded by the Wellcome Trust (Discovery Award 225880/Z/22/Z) or UK Medical Research Council (MC_UU00038/9). E.R., H.S., A.B., R.H., A.T. and A.J.F are funded by a UKRI Future Leaders Fellowship (MR/T043482/1) and E.R. is funded by an MRC PhD DTP programme (MC_ST_00034 Studentships).

## Materials and Methods

### Generation of metabolically labelled THP-1 cells

To generate heavy and light growth media, we reconstituted RPMI 1640 Medium for SILAC (Thermo Scientific 88365) with Arginine 240 mg/L (1.137 mM), Lysine 40 mg/L (0.219 mM) and Proline 180 mg/L (1.566 mM) (Sigma). Heavy isotopes were (L-)Arginine HCl ^13^C_6_ ^15^N_4_ (i.e. +10 mass) (Cambridge Isotope Laboratories CNLM-539-H) and (L-)Lysine ^13^C_6_ ^15^N_2_ (i.e. +8 mass) (Cambridge Isotope Laboratories CNLM-291-H), 10% dialyzed FBS (Gibco A3382001), 100 U/mL penicillin, 100 µg/mL streptomycin (ThermoFisher). Cells were passaged for 3 weeks, checking growth rates were comparable between the two lines. To test isotope incorporation, 3 million cells of the following conditions were lysed in 200 μL 8 M urea: light only, heavy only, light and heavy mixed 1:1 before lysis, light and heavy missed 1:1 after lysis. Lysates were vortexed, pelleted 12,000 xg 10 min, and 150 μL supernatant removed. 50 μg protein was made to 80 μL in 8 M Urea, to which 320 μL water was added, and DTT to 1 mM final concentration. Tubes were incubated 37 °C 30 min, cooled, then iodoacetamide (IAA) added to 5 mM final, and stored in the dark for 20 min. DTT was added to 5 mM final to quench unreacted IAA. TEAB pH 8 was added to 50 mM, reducing Urea to 1.5 M. Trypsin was added at 1:100 enzyme:substrate, and incubated 37 °C overnight. TFA 1% was added to samples and peptides were de-salted using c18 columns (Nest Group Microspin C18 HEM S18V), activated with 0.1% TFA, 100% acetonitrile. Peptides were eluted in 75% acetonitrile/0.1% TFA, frozen, and centrifuged to dryness. R+10, K+8 incorporation in the heavy line was 98.6 % at the time of experimentation.

### SILAC-ABPP of IFN treated THP1 cells

Eight T175 flasks each of Heavy or Light THP-1 were pooled, counted, and seeded in three 15 cm dishes, with Heavy or Light RPMI media, 50 million cells/dish, in the presence of 100 ng/mL PMA. This cell number provides ∼ 10 mg protein lysate from each cell line. 24 h later, media was refreshed. 2 d later, 1000 U/mL IFNα1 was added to all 3 dishes of Heavy THP-1; an equivalent volume of water was added to the Light THP-1. 24 h later, 10 μM MG132 was added to all 6 dishes for 4 h. Cells were harvested by scraping in ice cold PBS. Cells were washed thrice, and 3 dishes of Heavy or Light pooled and frozen. Each pellet was thawed on ice, lysed in 2 mL of ice-cold, degassed lysis buffer (50 mM Tris-HCl pH 7.5, 10 mM sodium 2-glycerophosphate, 50 mM sodium fluoride, 5 mM sodium pyrophosphate, 1 mM sodium orthovanadate, 270 mM sucrose, 50 mM sodium chloride, 200 μM phenylmethane sulfonyl fluoride (PMSF), 1 mM benzamidine, 10 μM TCEP, 1% NP-40). Incubated on ice 10 min, vortexed 10 s, centrifuged 17,000 rpm, 4°C, 30 min, and clarified lysates transferred to fresh tubes. Concentrations of each line were normalised to 4.5 mg/mL. 1 mL of each lysate was pre-cleared with 25 µL 50% equilibrated Streptavidin HiLink Ultra resin (Thermo Fisher) by rotation 4°C, 1 h. 0.5 mg of pre-cleared Heavy and Light lysates were combined to give 1 mg total protein. Biotin-UBE2D3∼Ub ABP was added at 3 μM final and reactions incubated at 30 °C, 4 h. 40 μL 50% equilibrated Streptavidin HiLink Ultra was added, and rotated 4 °C, 4 h. Parallel incubations were carried out with Biotin-UBE2L3∼Ub ABP and without ABP. Beads were then washed twice with 0.2 % SDS/PBS, twice with PBS, twice with 4 M Urea/PBS, twice with 1.5 M Urea/PBS, 50 mM TEAB (pH 8.0). All washes were performed by rotating tubes for 5 min. Beads were resuspended in 190 µL 1.5 M Urea, 50 mM TEAB (pH 8.0), to which 10 µL 100 mM TCEP added for a final concentration of 5 mM, incubated 56 °C, 30 min. 22 µL 100 mM IAA was added for a final concentration of 10 mM, incubated in the dark, 30 min. 25 µL 100 mM DTT added to final concentration 10 mM to quench IAA. 53 µL 50 mM TEAB (pH 8.0) added for final volume of 300 µL. 2 μg trypsin was added and incubate 37°C overnight. Samples were acidified with 1% TFA and cleaned as above.

LC–MS/MS analysis was performed on an LTQ Orbitrap Velos instrument (Thermo Scientific) with a Pepmap RSLC C18, 2 µm, 100 Å, 75 µM x 50 cm column (Thermo Scientific) coupled to an Ultimate Nanoflow HPLC system (Dionex). A gradient running from 3% solvent B to 99% solvent B over 345 min was applied (solvent A, 0.1% formic acid and 3% DMSO in H_2_O; solvent B, 0.08% formic acid and 3% DMSO in 80% MeCN). An inclusion list representing 5 tryptic peptides from each of the 42 HECT and RBR E3s was applied. An exclusion list representing the 5 major contaminating streptavidin peptides – identified from historic data – was also included. Differential signals were quantified by two channel processing (light vs. Arg10/Lys8) of the raw data file with Maxquant (version 1.6.5.0). Variable modifications included methione oxidation and N-terminal acetylation whereas carbamidomethylation of cysteine was considered a fixed modification. The human Swiss-Prot protein database (UP000005640) was searched with a peptide and protein false discovery rate of 0.01 and minimum peptide length of 7. Quantified protein groups with less than 1 unique peptide were manually excluded. The log2 transform of normalized H/L ratios was then plotted against protein ID in Prism (GraphPad).

### RT-qPCR validation of SILAC THP-1 cell lines

To test the IFNα1 response of THP-1 Light and Heavy cell lines, 50 thousand cells were stimulated with 1000 U/mL IFNα1 or water overnight. Cells were peletted and RNA extracted with a Qiagen RNeasy kit. 5 µL RNA was used for cDNA synthesis, and 2 µL cDNA was used in TaqMan qPCRs using 20X primer/probe mixes: ACTB (Hs01060665_g1); OAS1 (Hs00973635_m1); ISG15 (Hs00192713_m1)

### Expression of ZNFX1 in insect cells

Constructs harbouring C-terminally His tagged WT ZNFX1 (ZNFX1-His) or mutants (K625A, F1210A/R1216A, F1575A, Q1623A, C1860A, C1860K and H1881N) were cloned into pFastBacDual plasmids and transformed into DH10Bac *E. coli* cells. Blue/white screening selected for colonies successfully producing recombinant baculoviral shuttle vector (bacmids), which were purified by alkaline lysis and phenol/chloroform precipitation. Isolated bacmids were transfected into adherent Sf9 insect cells in 6 cm dishes using Cellfectin™ II Transfection Reagent (Thermo Fisher #10362100) and incubated for 96 hours at 27 °C to produce a low titer viral stock (P1) from which high-titer baculovirus was prepared following two rounds of transduction in adherent Sf9 cells (P2 and P3). Recombinant ZNFX1 was then expressed in a large suspension culture of Sf9 cells at 27 °C for 72 hours in un-supplemented Sf-900 II media (Thermo Fisher #10902104). Cells were harvested by centrifugation and snap-frozen in liquid nitrogen for storage at -80 °C. Cells were then lysed in lysis buffer (50 mM Tris-HCl pH 7.5, 150 mM NaCl, 25 mM Imidazole, 0.1 mM TCEP) supplemented with 50 μg/mL DNase I (Sigma Aldrich #DN25-100MG) and 1 x cOmplete EDTA-free protease inhibitor cocktail (Roche #5056489001) and subjected to brief sonication, before centrifugation at 40,000 x g and incubation with Nickel-NTA agarose resin (Qiagen # 30210). Resin was washed with wash buffer (50 mM Tris-HCl pH 7.5, 150 mM NaCl, 25 mM Imidazole, 0.1 mM TCEP) and eluted in equivalent buffer containing 300 mM imidazole. Protein was concentrated using a 100 kDa MWCO centrifugal filter (Millipore #UFC500324) and buffer exchanged into storage buffer (50 mM Tris-HCl pH 7.5, 150 mM NaCl, 0.1 mM TCEP) for long term storage at -80 °C.

### E2 expression and purification

All E2s employed in this study (UBE2L3 WT, UBE2D3 WT and mutants S22R, N77S, D87A, S94K, D117A, S22R/C85S, S22R/C85K, and UBE2L3) were expressed with N-terminal His-tags in BL21 *E. coli* cells and purified by nickel affinity chromatography as previously described^25^. Additional E2s employed in the E2 panel screen were produced by and purchased from MRC Reagents and Services, as were lysine-arginine Ub mutants.

### Expression and purification of ZNFX1 fragments

Sequences for human ZNFX1 covering unannotated regions 1462-1652 and 1533-1631 respectively were cloned into pGEX-6P-1 vectors, with the described mutations introduced by site-directed mutagenesis. WT ZNFX1_1462-1652_ and mutants F1575A and Q1623A, and ZNFX1^_^WT and F1575N were expressed as GST fusions in BL21 cells at 16 °C in media supplemented with 200 µM ZnCl2 and 100 µg/mL ampicillin. Proteins were purified by glutathione-sepharose chromatography, with ZNFX1_1462-1652_ WT and mutants eluted with 10 mM glutathione to retain the GST-tag (GST-ZNFX1_1462-1652_ WT, F1575A and Q1623A) whilst ZNFX1^1533–1631^ WT and F1575N underwent proteolytic cleavage on resin to produce the tag-free species. An additional construct harbouring His-SUMO tagged ZNFX1_1462-1652_ was also generated and expressed as a His-SUMO fusion as for the GST-tagged species and purified over Nickel-NTA agarose resin. Proteins were further purified by size exclusion chromatography over a Superdex 75 16/600 column (GE Healthcare), concentrated by centrifugal filtration and stored at -80 °C in 50 mM Tris-HCl pH 7.5, 150 mM NaCl, 0.1 mM TCEP.

### Multiple-Turnover Autoubiquitination Assays

All autoubiquitylation assays, including ubiquitin-mutant panels, were performed in 40 mM Tris-HCl pH 7.5, 150 mM NaCl, 10 mM MgCl2 diluted from a 10X stock and supplemented with 0.6 mM DTT and 4 mM ATP. E2 was preloaded with ubiquitin at 3X final enzyme concentration (1.25 µM E1, 12.5 µM E2, 125 µM ubiquitin) for 20 mins at 37 °C, before diluting to 1X with E3 ligase in reaction buffer, giving final concentrations of 0.4 µM E1, 4 µM E2, 40 µM ubiquitin and 1 µM E3. Reactions were then run at 37 °C for the indicated time, with isopeptide-trapping of Ub on ZNFX1 C1860K also achieved using this method. Where Cy3 or Cy5 labelled ubiquitin was used, the final ubiquitin concentration was 4 µM. Reactions were incubated at 37 °C for the timepoints indicated and quenched with 4X lithium dodecyl sulphate (LDS) + 715 mM β-mercaptoethanol (βME) to a final concentration of 1X or 2X except for E2 panel assays which were quenched only with LDS. Where indicated, E3 was supplemented with 10 µg/mL nucleic acid immediately before the addition of preloaded E2 mix, or pre-treated with RNase A (Thermo Fisher, #EN0531) at 100 µg/mL for 15 minutes at 37 °C. RNase A (EN0531; Thermo) in 50 % glycerol was prepared by 1/10 dilution in reaction buffer, with buffered 5 % glycerol serving as a control. For deubiquitinase (DUB), hydroxylamine (NH_2_OH) or sodium hydroxide (NaOH) treatments, autoubiquitination was ceased by inhibition of E1 with a E1-specific inhibitor (SHN-2028 C1, 25 µM)^79^, before adding 5 µM DUB, hydroxylamine (0.5 M, pH 9) or NaOH (0.14 M) for 30 mins at 37 °C. Reactions were then resolved by SDS-PAGE and visualised by either Coomassie staining, western blotting, or imaging by Chemidoc (Bio-Rad) using the appropriate channels for Cy3, Cy5, Fluorescein, Rhodamine or Coomassie detection (emission: 602/50 nm, 700/50 nm, 532/28 nm, 602/50 nm and 715/30 nm, respectively).

### Single-Turnover E2∼Ub Discharge Assays

E2 (10 µM), E1 (0.5 µM) and Cy3 labelled ubiquitin (12.5 µM) were incubated for 20 mins at 37 °C in buffer containing 40 mM Tris-HCl pH 7.5, 150 mM NaCl, 10 mM MgCl2 and 5 mM ATP. The loaded E2∼Ub reaction was cooled on ice for 2 minutes before addition of E1 inhibitor (SHN-2028 C1, 25 µM) for 10 mins at RT. An equal volume of E3 in reaction buffer was added, producing final enzyme concentrations of 5 µM E2, 0.25 µM E1, 6.25 µM ubiquitin and 0.2 µM E3. In specific assays where these concentrations are varied, the alternative concentration is indicated in the figure. Samples were quenched by 4 X LDS (Thermo Fisher), resolved by SDS-PAGE and visualised by Chemidoc (Bio-Rad) using the appropriate filter for Cy3 detection (602/50 nm). Data analysis was performed by band quantification on ImageJ (Fiji) and statistical analysis on Prism (GraphPad).

### Generation of stable E2∼Ub conjugates

UBE2D3 S22R variants with native Cys85, C85S, or C85K were used to prepare stable E2∼Ub conjugates with thioester, oxyester or isopeptide-linked ubiquitin respectively. Oxyester and isopetide (C85S and C85K) linked E2∼Ub were produced as previously described^38^; briefly 200 μM of E2 was incubated with 200 μM ubiquitin (Sigma, U6253-25MG) and 1 μM E1 at 37 °C overnight at pH 8.5 (C85S) or pH 10 (C85K) before purification by size-exclusion chromatography (SEC) and storage in 50 mM Tris pH 7.0,150 mM NaCl, 0.8 mM TCEP. For the thioester, a large-scale loading reaction was conducted as described for multiple- and single-turnover autoubiquitination assays. Loaded E2∼Ub was then purified by SEC and stored in 50 mM Na_2_HPO_4_ pH 6.0, 150 mM NaCl without reducing agent at -80 °C.

### Quantitative ATPase Assay

ATPase activity of WT and mutant ZNFX1 was assessed using the Malachite Green Phosphate Detection Kit from R&D Systems (DY996) following the standard protocol. Reactions were conducted in 50 mM Tris, 150 mM NaCl, 5 mM βME, 40 mM MgCl_2_, and initiated by adding concentrated ZNFX1 (16X) to ATP (20 µM) to give final concentrations of 0, 12.5, 25 or 50 nM ATPase. After the appropriate time (up to 160 mins), reactions were quenched with 20 µL Malachite Green working reagent and incubated for a further 30 mins RT before absorbance measured at 620 nm on a BioTek Epoch Microplate Reader. Orthophosphate standards and quenching reagent were prepared fresh each time in accordance with the standard protocol. Data was subsequently analysed using Microsoft Excel and GraphPad Prism.

### Isothermal Titration Calorimetry (ITC)

ITC experiments were conducted as previously described^60^ using a MicroCal PEAQ-ITC instrument with PEAQ-ITC Software (ver. 1.40) (Malvern Panalytical). UBE2D3 was added to the cell at 94 µM concentration, with ZNFX1_1533-1631_ WT or F1575N in the syringe at 1300 µM. An initial dummy injection of 0.4 µL was followed by 12 injections of 3.5 µL, and a reference trace of ZNFX1 titrated into buffer-only was deducted to normalise the titrant dilution enthalpy. Reactions were conducted at 25 °C and analysed using the PEAQ-ITC software.

### Activity-Based Probe (ABP) Labelling

Full-length ZNFX1 was captured using a biotinylated E2∼Ub activity-based probe (ABP) with a cysteine-electrophile trap^25^ for cross-linked (XL) peptide mapping, whilst ZNFX1_1462-1652_ was assessed for allosteric ligase activity using a photocrosslinking E2∼Ub ABP^57^. 35 μg WT ZNFX1 was incubated with 175 μg biotinylated UBE2D2∼Ub probe (8 molar eqs.) in 50 mM Na_2_PO_4_ pH 7.5, 150 mM NaCl in a final volume of 250 μL for 4 hours at 30 °C. Labelling of ZNFX1 was determined by SDS-PAGE and Coomassie stain, with labelled and unlabelled ZNFX1 subsequently separated from unreacted probe by gel filtration over a Superdex 200 pg Increase 10/300 GL column (GE healthcare). Fractions were pooled and concentrated by centrifugal filtration (Millipore #UFC500324), before incubation with pre-equilibrated streptavidin resin (Thermo Fisher #88816) for 3 hours at RT. Beads were washed, reduced and alkylated as previously described ^25^, and resin-bound proteins were digested with GluC and trypsin proteases overnight. Resultant peptides were desalted and freeze-dried prior to resuspension in 5% formic acid in water for LC-MS/MS analysis. 2 μg of peptides were injected on an UltiMate 3000 RSLC nano System coupled to an Orbitrap Fusion Lumos Tribrid Mass Spectrometer (Thermo Fisher Scientific). Peptides were loaded on an Acclaim Pepmap trap column (Thermo Fisher Scientific #164564-CMD) prior to analysis on a PepMap RSLC C18 analytical column (Thermo Fisher Scientific #ES903) and eluted on a 60 or 120 min linear gradient from 3 to 60% Buffer B (Buffer A: 0.1% formic acid in water, Buffer B: 0.08% formic acid in 80:20 acetonitrile:water (v:v)). Eluted peptides were then analysed by the mass spectrometer operating in data dependant acquisition mode using a TOP 3s method.

Raw LC-MS/MS data was searched for cross-linked (XL) peptides using pLink (Chen, Meng et al. 2019) and searched against a reference sequence catalogue containing sequences for *Hs*ZNFX1, *Hs*UBE2D2* and *Hs*Ubiquitin (Uniprot IDs Q9P2E3, P62837 and P0CG47 respectively; UBE2D2* harbouring C21S C107S C111S mutations and Ub sequence 1-73 only). A reverse sequence database was generated to determine a false discovery rate (FDR), with peptides filtered to 1% FDR. A crosslinker mass of 306.181 Da and elemental composition of C(14)H(22)N(6)O(2) was entered manually, and the following variable modifications were included: carbamidomethyl cysteine, methionine oxidation/dioxidation, asparagine and glutamine deamidation. Minimum peptide length was established at 6 residues, maximum peptide mass 10 kDa with 3 missed cleavages and peptide and fragment mass tolerance set at plus or minus 10 ppm. Scans identified as containing a relevant XL-peptide spectra were analysed using pLabel to assess fragmentation and ion abundance.

The photocrosslinking ABP was generated and applied as previously described ^57^. Briefly, labelling reactions were performed with 10 μM ZNFX1_1462-1652_ (WT or F1575A) and 20 μM ABP harbouring WT or F62A UBE2D3. An RNF4 fused dimer was employed as a positive control. Reactions (35 μL) were conducted in a 24 well plate in 20 mM Hepes, pH 7.5, 150 mM NaCl, and 1 mM TCEP. Samples were then split, with one half irradiated with UV light (365 nm) for up to 30 minutes whilst the remaining portion was maintained in the dark. Reactions were conducted on ice at 10 cm from the UV light source and quenched with addition of 4X LDS+βME and resolved by SDS-PAGE and visualized by Coomassie stain. Timepoints were periodically collected from a single well.

### Computational Modelling with AlphaFold2

AlphaFold models of Ub:UBE2D2:ZNFX1_1430-end_ and Ub:UBE2D2:ZNFX1_1430-1843_ complexes were produced using ColabFold (AlphaFold2 + MMSeqs2) using default settings^80^. All generated models for each complex produced almost indistinguishable structures, with the ‘Best Model’ as specified by AlphaFold used for structural analysis of each complex. Structures were examined using ChimeraX 1.4)^81^, and structural alignments were performed using the ChimeraX ‘matchmaker’ tool which superposes structures based on pairwise sequence alignment and individual fitting of residue pairs. Structures used for alignments include MYCBP2 and Upf1^25,64^.

### Sequence Alignments

Sequences were aligned using the Clustal Omega web service. Alignments of more than three sequences were analysed using the multiple sequence alignment tool (MSA), whilst alignments of only two sequences were performed using the pairwise alignment tool (PA) using the Needleman-Wunsch algorithm for global alignment (EMBOSS Needle). Alignments were further analysed using Jalview and annotated in Adobe Illustrator.

### Structure-guided sequence alignment using PROMALS3D

Alphafold models of the U-box domains from the selected E3 ligases were generated using UniProt-defined domain boundaries. For *Sc*Prp19p, the experimental structure was used (PDB: 2BAY). Structural alignments were performed in UCSF ChimeraX using the MatchMaker tool. Key residues within the hydrophobic core and hydrogen bond (H-bond) network of the U-box fold were manually inspected across models ^82^. Based on these comparisons, alignment constraints were applied to promote the correct positioning of structurally conserved residues. Final multiple sequence alignment was performed using PROMALS3D, which integrates sequence information with predicted secondary structure and 3D structural data to improve alignment accuracy.

### In-Vitro Transcription (IVT)

Single-stranded RNA (ssRNA) was generated using the HiScribe® T7 High Yield RNA Synthesis Kit (E2040S; NEB) following the standard protocol. For the template, genes for AMPKA2 (*PRKAA2*), AMPKB2 (*PRKAB2*) and AMPKG1 (*PRKAG1*) were cloned sequentially into a pOPC Lac vector harbouring a T7 RNA polymerase promoter upstream of the three sequences and a T7 terminator sequence downstream. The plasmid was linearised by restriction digestion, with cleavage upstream of the T7 promoter by PshAI producing a 4 kbp template for transcription, running from T7 promoter to terminator. Alternatively, the plasmid was linearised with BamHI downstream of the T7 promoter, between *PRKAA2* and *PRKAB2* genes, producing a 1.8 kbp template from T7 promoter to the cleavage site. Linearised plasmid was purified by agarose gel extraction (T1020S; NEB) or using the Monarch DNA cleanup kit (T1030; NEB) and incubated with T7 RNA polymerase and free RNA nucleotides (ATP, GTP, CTP and UTP) following the HiScribe® kit protocol. Resultant RNA from the two *PRKAA2* templates and the provided Fluc Control Template (1.8 kb) was purified using the Monarch RNA cleanup kit (T2030S; NEB) and quantified by nanodrop, producing three ssRNA species: *PRKAA2-PRKAB2-PRKAG1* mRNA in tandem (ssRNA*_PRKAA2-L_*), *PRKAA2* mRNA only (ssRNA*_PRKAA2-S_*), and the Fluc control mRNA (ssRNA_Fluc_). All other nucleic acids used in the study were purchased from Invivogen, including: high-molecular weight poly(I:C) (tlrl-pic), low molecular weight poly(I:C) (#tlrl-picw), fluorescein-labelled poly(I:C) (#tlrl-picf), rhodamine-labelled poly(I:C) (#tlrl-picr), biotin-labelled poly(I:C) (#tlrl-picb), and poly(dA:dT) (#tlrl-patn).

### Western Blotting

Proteins were separated by SDS-PAGE on 4-12 % Bis-Tris precast polyacrylamide gels (NuPAGE™) in 2-(N-morpholino)ethanesulfonic acid (MES) at 200 V, before transfer to polyvinylidene fluoride (PVDF) membrane at 85 volts for 90 minutes in Tris-glycine (48 mM Tris, 39 mM glycine, 20 % MeOH). Membranes were incubated for 1 hour in either 5 % bovine-serum albumin (BSA) (Sigma; A7906) or milk powder (Marvel) (w/v) in tris-buffered saline with 0.1 % TWEEN-20 (Sigma; P1379-100ML) (TBST), before addition of primary antibodies (anti-ubiquitin pAb, Z0458, DAKO; anti-ZNFX1 mAb, ab179452, Abcam,or Abcam EPR12330; anti-Ub-K63 HWA4C4 mAb, 14-6077-82, eBioscience^TM^/Thermo; anti-Ub K48 Apu2 ZooMAb®, ZRB2150-4X25UL, Sigma; 6xHis mAb, 631212, Clontech, anti-Tubulin (1:2000, Cell Signalling Technology, 21445), anti-Vinculin (1:2000, Cell Signalling Technology, 139015), anti-SINV capsid (1:2000 ^35^), anti-puromycin (1:4000, Millipore, MABE343), anti-GAPDH (1:5000, Invitrogen, AM4300), anti-actin (1:5000, Cell Signalling Technology, 3700S)), for 1 hour at RT or 4 ^°^C overnight. Membranes were washed in TBST before addition of the appropriate HRP-conjugated secondary antibody (1 h RT) (anti-mouse IgG HRP, 7076S, CST; anti-rabbit IgG HRP, 7074S, CST), before further washing with TBST and visualisation using enhanced chemiluminescence (ECL) Western blotting substrate (Pierce; 32106) and visualised by ChemiDoc™ (Bio-Rad) imaging system, a LICOR Odyssey XF Scanner, or exposed to radiographic film on an SRX-101A Medical Film Processor (Konica Minolta). Bands were quantified using Image J, normalising to the background signal and the housekeeping gene.

### Semi-Denaturing Detergent Agarose Gel Electrophoresis (SDD-AGE)

SDD-AGE was performed as described^68^. ZNFX1 was driven to aggregate via multiple-turnover autoubiquitination assay in the presence of RNA and ATP and reactions quenched with 2X LDS sample buffer without βME and boiled 5 mins 95 °C. Samples were resolved by 1.5 % agarose / 0.1 % SDS gel in 1 X TAE/ 0.1 % SDS buffer at 100 V, monitoring periodically for the progression of pre-stained protein ladder (1610373; Bio-Rad). Gels were stained overnight in Coomassie Instant Blue (ab119211; Abcam), and destained in water before visualisation by Coomassie far-red detection (715/30 nm) ChemiDoc™ (Bio-Rad) imaging system.

### Ub-Chain Induced Aggregation

ZNFX1 with or without RNA and/or ATP, free Ub or free Ub-chains, and E2∼Ub thioester were each prepared as three 3X stocks in 40 mM Tris-HCl pH 7.5, 150 mM NaCl, 10 mM MgCl2 and mixed 1:1:1, giving final concentrations of 1 µM ZNFX1, 10 µg/mL RNA (ssRNA_Fluc_), 3 µM ATP, 5 µM (low) or 25 µM (high) free Ub, or 10 µM of either K63-linked or K48-linked tetra-Ub (MRC Reagents and Services), and E2∼Ub thioester at 4 µM. Reactions were incubated for 1 hour at 37 °C and visualised by SDS-PAGE, Coomassie stain and Coomassie far-red detection (715/30 nm) with the ChemiDoc™ (Bio-Rad) imaging system.

### Liquid Chromatography - Mass spectrometry measurement RNA nucleotide ubiquitination

ZNFX1 conjugation reactions containing E1 (0.4 µM), UBE2D3 (4 M), Ub (40 µM) ZNFX1 (1 µM) and either no nucleotide or no ATP were performed. The nucleotides AMP, CMP, GMP and UMP were added at 38 mM, which approximates that of model small molecule substrates modified by E3 ligases. Reactions were incubated at 37 °C for 1 h and resolved with a 10-70% gradient over 20 minutes (mobile phase A: H2O + 0.05% TFA; mobile phase B: acetonitrile + 0.04% TFA) using an Agilent Affinity 1200 single quadrupole LCMS system. Deconvoluted protein masses were obtained using the Chemstation software (Agilent).

### Native PAGE

T-REx-293 cells were treated with 1 μg/mL tetracycline for 24 h, and where indicated, either harvested or infected with SINV-WT at an MOI of 1 for 1, 3, 5, or 15 h before harvest. Cells were washed and lysed in 1X NativePAGE buffer with 1% DDM at 4 °C (Life Technologies). The lysates were normalised, and the G-250 dye added to a final concentration of 0.25%. Native gel electrophoresis was performed using the Life Technologies NativePAGE Novex Bis-Tris gel system. Samples were kept on ice throughout, until being loaded into the gel. Electrophoresis was performed with the tank in an ice bath. The proteins were transferred to a PVDF membrane and stained with antibody as described above. Where indicated, poly(I:C) (Invivogen, tlrl-pic), or a de-ubiquitinase, GST-USP2 (MRC Reagents and Services), was added directly to the lysate, before the addition of G-250 dye.

### Live Cell Microscopy

T-REx-293^ZNFX1^ ^KO^ cells reconstituted with mNeonGreen-ZNFX1 were seeded at 6 x10^5^ in 30 mm glass-bottomed dishes (Thermo Scientific) with 1 μg/ml tetracycline and incubated for 24 h. Fluorobrite DMEM (Gibco) supplemented with 10% FBS and 1 μg/ml tetracycline, containing SINV-mCherry for an MOI of 20, was added to the cells. The dish was placed in the heated chamber of the microscope, warmed to 37°C and with CO_2_ at 5%. Images were taken every 10 min in the GFP and mCherry channels. All images were collected with a Axio Observer.Z1 microscope equipped with Plan-Apochromat 63x/1.40 Oil DIC M27. Definite Focus was used for maintenance of focus over time. Images were acquired with a Axiocam 506 imaging device controlled with ZEN software. ImageJ software was used to remove shot pixels with the remove outliers tool. Imaris Software was then used to perform deconvolution on both channels. ImageJ was then used to perform bleaching correction on the GFP channel, background subtraction on both channels, and alter the brightness and contrast.

### Cell proliferation assay

T-REx-293 cells were seeded in duplicate at a density of 2.4 x 10^5^ in 24 well plates in DMEM supplemented with 2% FBS and 1 μg/mL tetracycline. At 24, 48, 72, and 96 hours, the supernatant was removed and 100 µL of trypsin was added. The cells were incubated at 37 °C for 5 min followed by the addition of 900 μL DMEM. The cells were mixed thoroughly and diluted 1:1 in Trypan Blue. The cell concentration was measured on a Countess cell counter (Invitrogen), with both total and live counts recorded.

### Generation of ZNFX1 KO cell lines

To assess the effect of ZNFX1 on SINV replication a ZNFX1-null cell line was generated in a T-REx™-293 Flp-In background. Oligonucleotides encoding ZNFX1-specific guide RNA sequences (guide 1: TCCTAACCAACGAGTCTGTT; guide 4: AGCGGTTCTGACAGGTCCGC) were annealed and cloned into px459 (pSpCas9(BB)-2A-Puro (px459) (a gift from Feng Zhang (Addgene plasmid # 48139; http://n2t.net/addgene:48139; RRID:Addgene_48139)). Sequence-verified plasmid was transfected into T-REx™-293 cells using polyethylenimine (PEI), transfectants were selected with 2.5 μg/mL puromycin 1 d later, then single-cell cloned by limiting dilution in 96 well plates 2 d after this, in the absence of puromycin. Five clones were screened by Western blot using anti-ZNFX1 (Abcam, EPR12330) and anti-β-Actin (Cell Signaling Technology, #3700). Genomic DNA was extracted from triaged clones (example of 3 clones in **Supplementary Fig. 2a**) using a DNeasy Blood & Tissue kit (Qiagen) and a PCR performed to amplify a region flanking the intended edit site (For guide 1: RH03 TGGCCATTTACTAATGAGCCTAT and RH04 GGGCCAGGTTGAATAAGCTC; for guide 4: RH07 GCAGATATCCAACCGCATCT and RH08 GCTTCCATCTTTTCCTGCAC). Amplified product was purified (Monarch PCR and DNA Cleanup Kit) and sequenced with a nested primer (for guide 1: RH01 TGGGTGACAGAGTGCAAGAG; for guide 4: RH05 GGACTGTGTGCAGCTGTGTT). Amplicons from the parental cell line were included as a control for the sequencing reaction. Clone 7 was chosen as the representative knockout clone for guide 1 and used in all reconstitution experiments described. Clone 3 from guide 4 transfected cells contained the homozygous edit resulting in ZNFX1 protein pT1489Rfs*53.

### Generation of ZNFX1 variants

ZNFX1 variants were generated by *in vivo assembly* as previously described ^29^ using the following primers for mutagenesis.

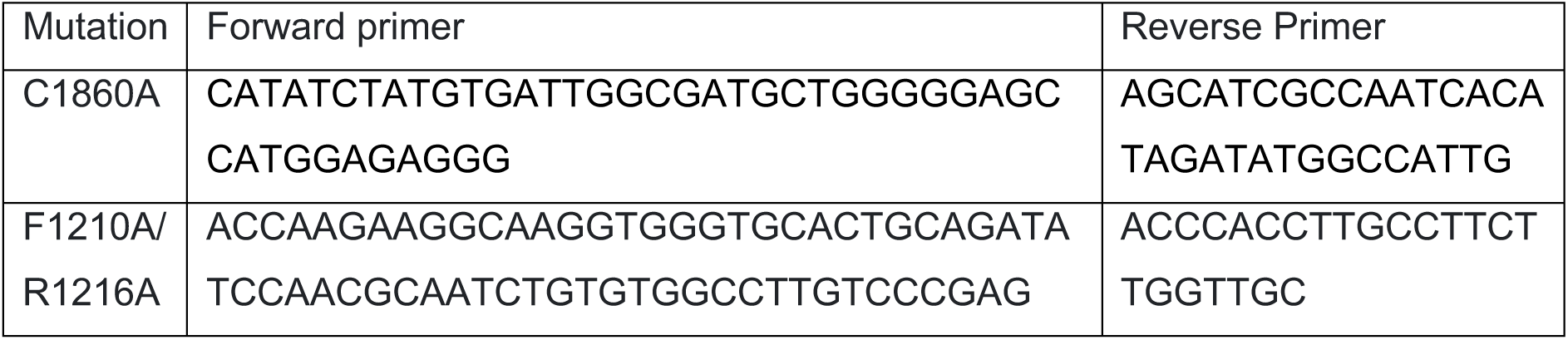
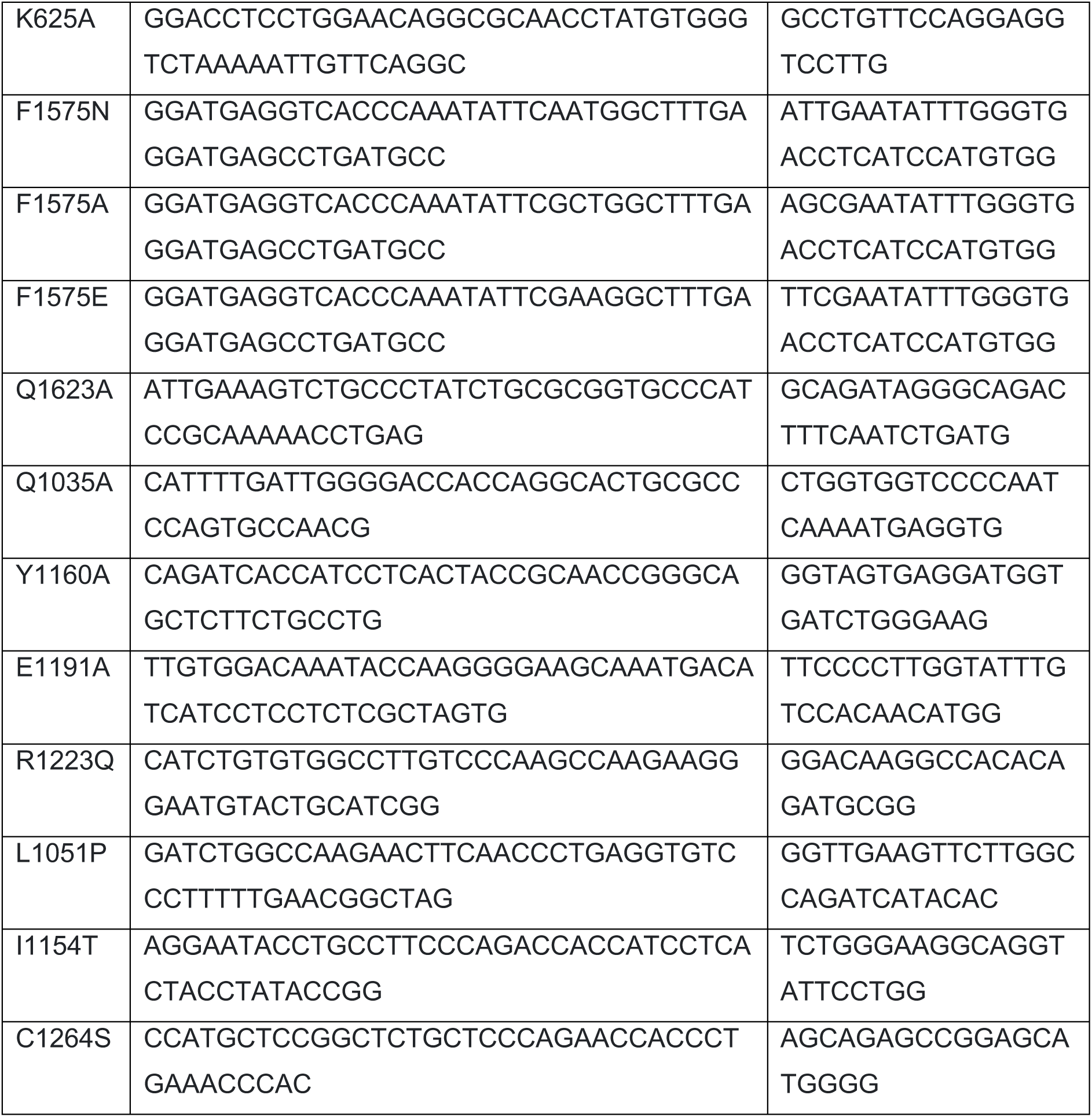

mNeonGreen-ZNFX1 was generated by Gibson Assembly following independent amplification of ZNFX1 and mNeonGreen, using the primers RH96 GGTGGTGGTATGGAGGAGAGA and RH97 to amplify ZNFX1, and primers RH98 AGGAGATCCAGGGGATGATGTAGCCCGGGGAATTCGCTAGC TCTCTCCTCCATACCACCACCCTTGTACAGCTCGTCCATGCC FRT/TO-mNeonGreen. to and amplify RH99 pcDNA5-

### IVT SINV production and propagation

pT7-SVwt, pT7-SVmScarlet, and pT7-SVmCherry were received as a kind gift from Alfredo Castello. Virus was rescued from the plasmid as described (Garcia-Moreno et al., 2019) propagated in BHK21 cells. Plaque assays to calculate infectious titres were performed as follows:

VeroE6 cells were seeded at 3x10^5^ cells/well in a 12-well plate and grown to confluency overnight. Cells were inoculated with 250 µL SINV serially diluted tenfold in serum-free DMEM and incubated at 37 °C for 1 h. An overlay of 1:1 Avicel (1.2%; FMC) and 2× MEM (Gibco), supplemented with 2% FBS was then added. Cells were incubated for two or three days, fixed with 10% formaldehyde solution, washed twice with PBS (Gibco), and stained with Coomassie Blue.

### Cell culture and maintenance

T-REx-293-Flp-In and VeroE6 cells were propagated (37 °C, 5% CO2) in Dulbecco’s Modified Eagle Medium (DMEM; ThermoFisher) supplemented with 10% (v/v) fetal bovine serum (FBS; ThermoFisher) and antibiotics (100 U/mL penicillin, 100 µg/mL streptomycin; ThermoFisher).

### Virus replication assays

T-REx-293 cells were seeded at a density of 4x10^4^ in each well of a 96-well plate in DMEM supplemented with 2% FBS and 1 μg/mL Tetracycline. They were infected with SINV-mCherry or SINV-mScarlet at 0.1 MOI in the same media, or with human metapneumovirus-GFP at a range of dilutions, in a final volume of 200 µL. Where indicated, cells were treated with Ruxolitinib at 2 μM in 20 µL, or a mock treatment with DMSO, for 2 h prior to infection. Cells were incubated at 37 °C and 5% CO_2_ in a CLARIOstar fluorescence plate reader (BMG Labtech) for over 30 h. Fluorescent signal was monitored by measuring fluorescence (mCherry/mScarlet: excitation 570 nm, emission 620 nm; GFP: excitation 470 nm, emission 515 nm) every 15 min.

### RT-qPCR

T-REx-293 cells were treated with 1 μg/mL Tetracycline and seeded at 4x10^5^ in 12 well plates in DMEM supplemented with 2% FBS and 100 U/mL penicillin, 100 µg/mL streptomycin (ThermoFisher) as previously described. After 24 hours cells were infected with WT SINV at an MOI of 0.1. At the indicated timepoints post-infection the supernatant was removed, and the cells were lysed in TRIzol (Thermo Fisher Scientific). Total RNA was then extracted from the TRIzol samples using phenol-chloroform extraction and RNA-easy columns, with an on-column DNase treatment (Qiagen). cDNA was synthesised using SuperScriptIII (Invitrogen, 56575) with random hexamer primers (Invitrogen, 100026484). The cDNA was then treated with RNaseH. Host and viral gene expression was measured using TaqMan Fast Universal Master Mix (Applied BioSystems) and specific TaqMan probes listed below on the QuantStudio 3 Real-Time PCR machine (Thermo Fisher Scientific). Using the 2^-△△Ct^ method, viral transcript levels were normalised to ACTB and then normalised to input viral transcripts at 2 h in the respective cell line.

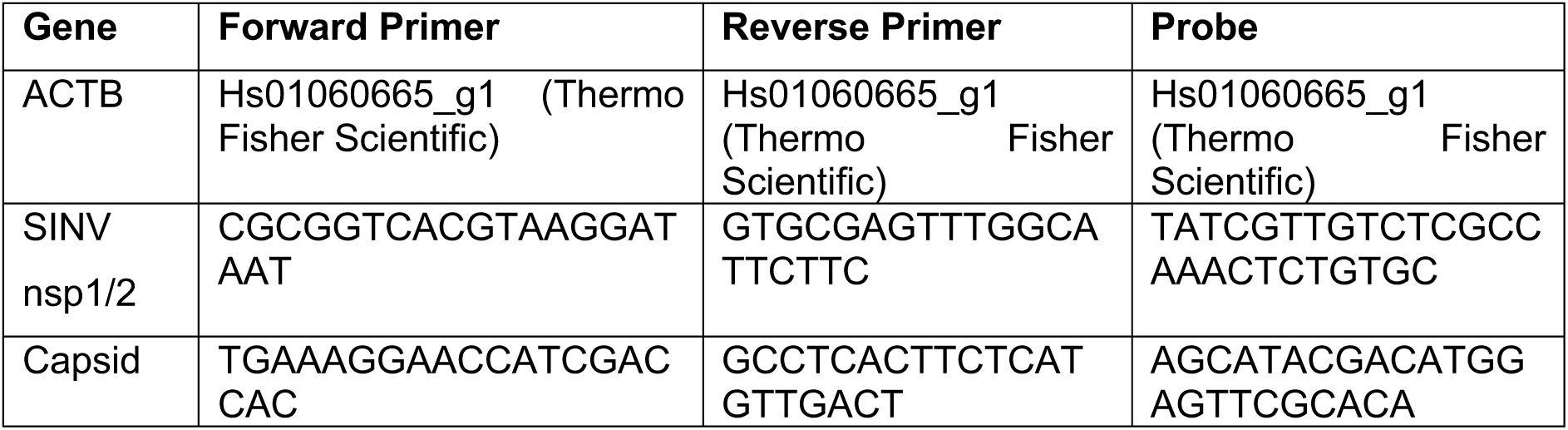

### Flow cytometry

T-REx-293 cells were seeded at a density of 4x10^4^ in 96 well plates in DMEM supplemented with 2% FBS and 1 μg/mL Tetracycline and then infected with mCherry-SINV or mScarlet-SINV at an MOI of 0.01. 24 h post-infection the cells were trypsinised and fixed in 4% paraformaldehyde. The percentage of mCherry or mScarlet positive cells was determined using flow cytometry (Guava EasyCyte, Merck) and calculated using FlowJo.

### Western blot for infection

T-REx-293 cells were treated with 1 μg/mL tetracycline and 4x10^5^ cells were seeded in each well of a 12 well plate in DMEM supplemented with 2% FBS. After 24 hours cells were infected with WT SINV at an MOI of 0.1. At 16 or 18 hours post infection the cells were washed, harvested, and lysed in lysis buffer. Clarified lysates were then normalised by Bradford assay and prepared for analysis by Western Blot in LDS sample buffer (Thermo Fisher Scientific) containing 2-mercaptoethanol. Where indicated, the cells were also treated with Bortezomib at 100 nM at the time of infection or puromycin was added to the cells to a final concentration of 5 ng/mL for 1 h prior to collection of the cells.

### Cycloheximide Chase

T-REx-293 cells were seeded at 6x10^5^ in each well of a 6 well plate and treated with 0.1 μg/mL Tetracycline. After 24 h the cells were treated with Cycloheximide (Merck, 0.1 mg/mL) and harvested at the indicated time after treatment. Where indicated, the cells were also treated with DMSO, 10 µM MG132, 100 nM Bafilomycin A1. The cells were lysed in lysis buffer and clarified lysates were normalised by Bradford assay and prepared for analysis by Western blot in LDS sample buffer (Thermo Fisher Scientific) containing 2-mercaptoethanol.

### Cellular poly(I:C) binding assay

T-REx-293 cells were treated with 1 μg/mL Tetracycline. After 24 h cells were harvested and lysed. The clarified lysates were normalised by Bradford assay and an input sample was taken. 5 μg of High Molecular Weight Biotin-Poly(I:C) (Invitrogen). The lysates were incubated for 1 h at 4°C with gentle agitation. Magnetic streptavidin beads (Pierce, Thermo Scientific) were equilibrated with lysis buffer and then 10 μL was added to each sample. After 1 h incubation at 4°C with gentle agitation, the beads were washed 4 times in lysis buffer and bound protein was eluted into 2x LDS containing BME by heating at 95 °C for 5 min. The input samples and eluted proteins were analysed by immunoblotting with anti-ZNFX1. For experiments using ATPγS, the indicated amount of ATPγS was added just prior to the addition of the biotin-poly(I:C). Additionally, the incubations were both shortened to 10 min each.

### Software, figure preparation (incl R)

Graphs were generated using R ^83^. Structural models were designed in Chimera X. Figures were prepared in Adobe Illustrator.

