## Supplemental figures for "ZNFX1 uses two-component ubiquitin circuitry to quarantine viral RNA"

**Supplementary Figures**

Daniel R. Squair[1,‡], Eilidh Rivers[2,‡], Hanna Sowar[2], Arda Balci[2], Roosa Harmo[2], David J. Wright[1], Gaurav Beniwal[1], Mathieu Soetens[1], Sunil Mathur[1], Aidan Tollervey[2], Callum Stanton[1], Adam J. Fletcher[2,*] and Satpal Virdee[1,*]

[1]MRC Protein Phosphorylation and Ubiquitylation Unit, School of Life Sciences, University of Dundee, Dundee, United Kingdom.

[2]MRC University of Glasgow Centre for Virus Research, University of Glasgow, Glasgow, United Kingdom

‡These authors contributed equally

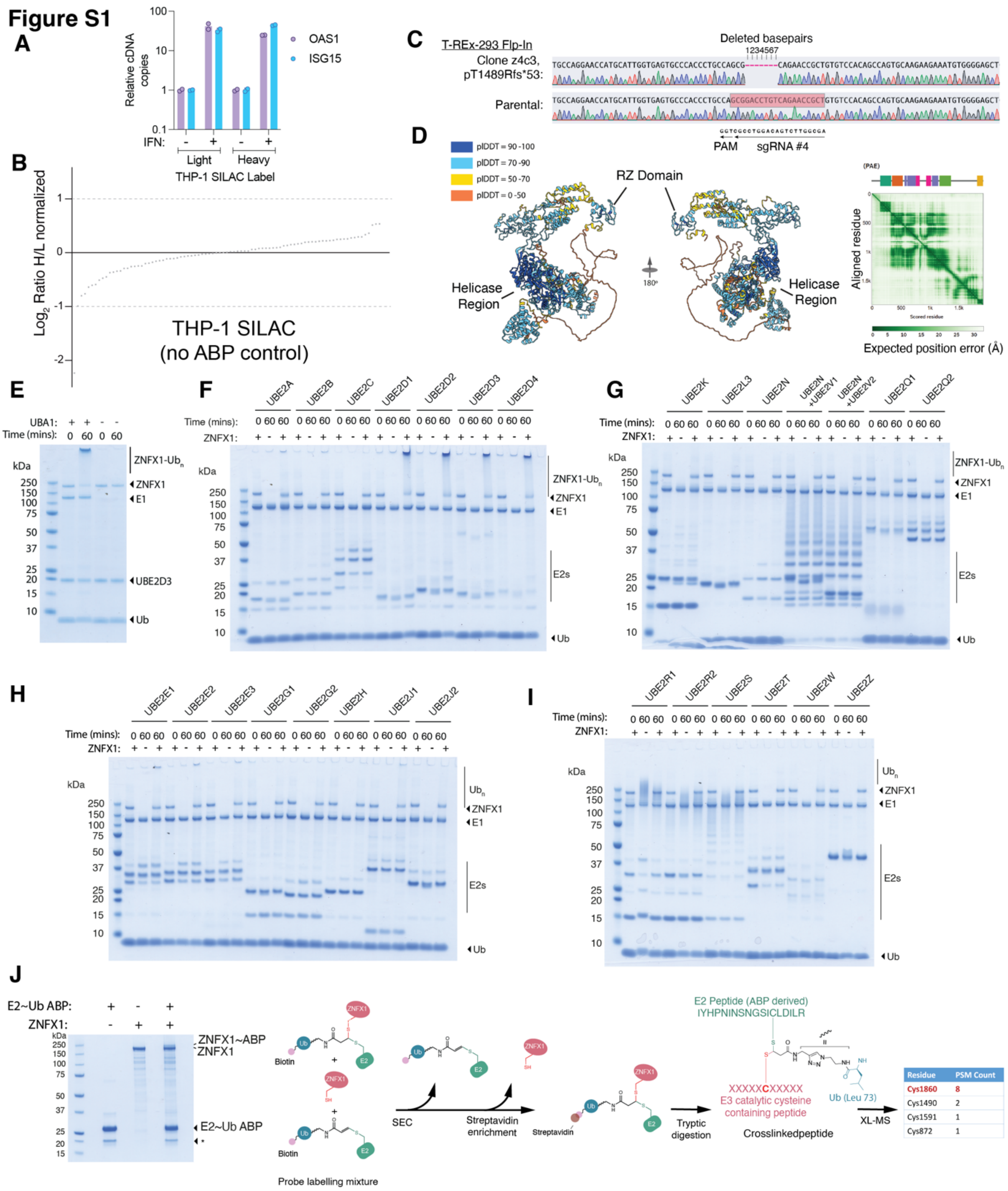


### Supplementary Figure 1

**A)** Metabolic labelling does not affect upregulation of the ISGs *OAS1* and *ISG15* in response to IFNα1, as determined by qPCR. 1000 U/mL IFNα1 or water was added to undifferentiated Heavy or Light THP-1 cells overnight and cDNA prepared the following day. **B)** Waterfall plot depicting the 87 quantitated SILAC ratios in the control experiment where ABP was withheld. No ratios were determined for any E1 activating enzymes or E3 ligases. **C)** Sanger sequencing analysis of the CRISPR/Cas9 edited T-REx-293 Flp-In clonal cell line (pT1489Rfs*53) which has deleted the last 429 residues. **D)** AlphaFold2 structural model of full-length human ZNFX1. The predicted model is coloured by pLDDT score, with annotated helicase and RZ domains indicated. Structural prediction was performed using ColabFold with default settings. Predicted Aligned Error (PAE) and identified Encyclopaedia of Domains (TED) is shown. **E)** ZNFX1 autoubiquitination assay in the presence and absence of E1 enzyme (UBA1). Formation of the high molecular weight smear is dependent on E1, consistent with ZNFX1 undergoing autoubiquitination. **F – I)** ZNFX1 (1 µM) was screened against a panel of E2 enzymes (4 µM) to identify those supporting ZNFX1-dependent autoubiquitination. Members of the UBE2D1-4 family showed the highest activity. Individual E2s and reaction times are indicated in the figure. Reactions were quenched with non-reducing 1X LDS sample buffer. **J)** ZNFX1 (3 µM) was incubated with biotinylated UBE2D3-Ub activity-based probe (25 µM) for 4 h at 37 °C. Unreacted probe was removed by size-exclusion chromatography (SEC), and the probe-labelled fraction of ZNFX1 was enriched using streptavidin resin. Bound material was subjected to on-resin tryptic digestion followed by crosslinking mass spectrometry (XL-MS). A schematic of the enrichment strategy and the crosslinked species is shown along with the identified peptide spectral matches (PSMs).

Figure S2

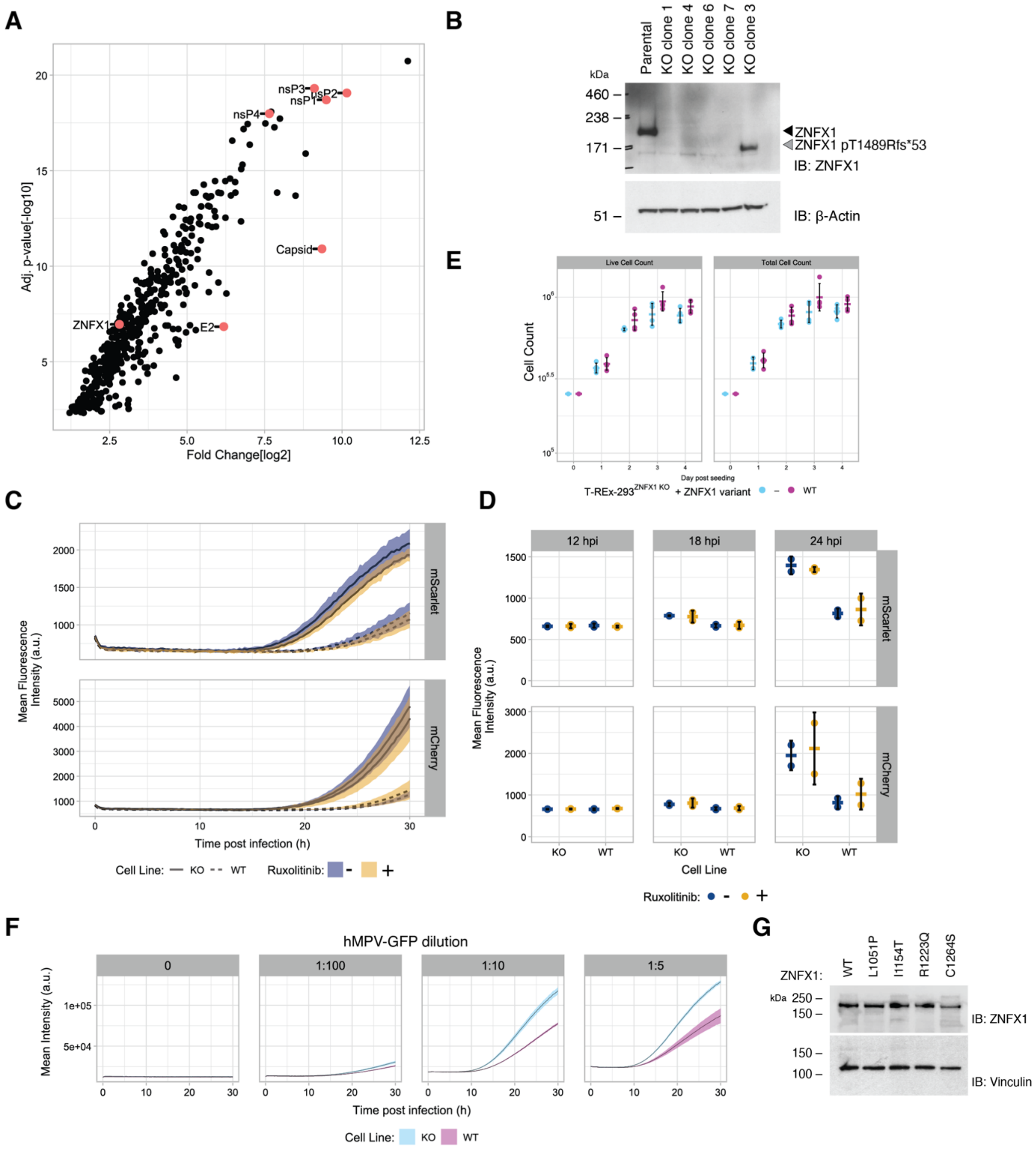


### Supplementary Figure 2

**A**) Published data from viral RNA interactome capture (vRIC) of SINV in HEK293 cells [1]. ZNFX1 is identified as an RNA binding protein recruited to SINV genomes. **B**) T-REx-293 Flp-In clones transfected with px459-expressing ZNFX1-specific sgRNAs 1 or 4, and Cas9, compared to the parental cell line. ZNFX1 expression determined by Western blot, β-actin served as a loading control. SgRNA1, clone 7, was used for subsequent experiments. **C**) Red fluorescent signal in T-REx-293$^{ZNFX1\ KO}$ cells (solid line) reconstituted with WT ZNFX1 (dashed line) infected with SINV-nsp3-mScarlet (top panel) or SINV-mCherry (bottom panel).

Fluorescence was measured at intervals of 15 min in an incubated plate reader (37°C and 5% $CO_2$). Data are presented as mean values ± SD from 3 independent infections. **D**) Mean ± SD of red fluorescent signal from two biological replicates of (C) at 12, 18, and 24 hours post infection. **E)** Cell proliferation of T-REx-293$^{ZNFX1\ KO}$ cells (blue) reconstituted with WT ZNFX1 (pink), was measured every 24 h by quantifying cell density. Wells were collected in duplicate and counted in duplicate, and mean ± SD is shown. **F)** Green fluorescent signal in T-REx-293$^{ZNFX1\ KO}$ cells (blue) reconstituted with WT ZNFX1 (pink) infected with human metapneumovirus-GFP. Fluorescence was measured at intervals of 15 min in an incubated plate reader (37°C and 5% $CO_2$). Data are presented as mean values ± SD from 3 independent infections. **G)** Expression levels of ZNFX1 variants indicated, in T-REx-293$^{ZNFX1\ KO}$ cells, was determined by Western blot. Vinculin served as a loading control.

**Figure S3**

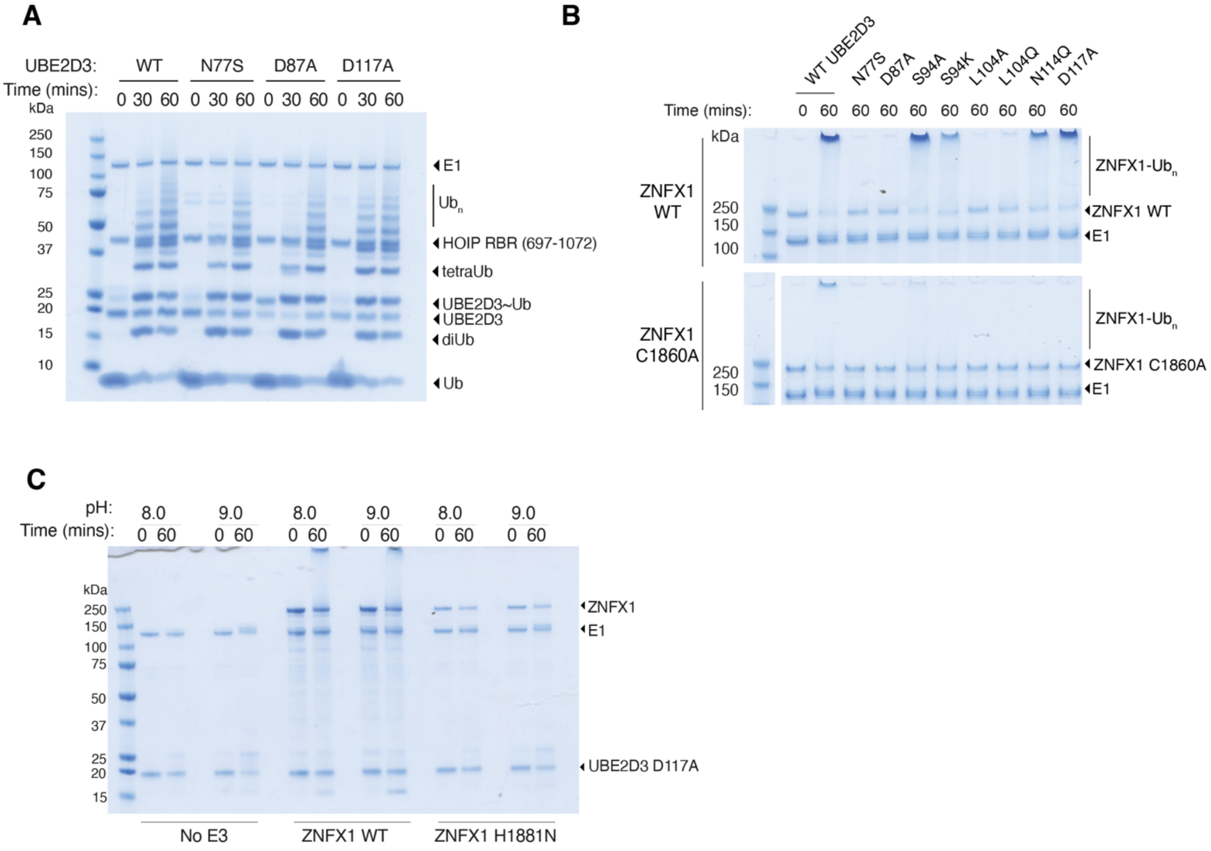


**Supplementary Figure 3**

**A)** Multiple-turnover ubiquitination assay with the catalytic domain of the RBR E3 HOIP (residues 697-1072; DU DU22629) and either wild-type (WT) UBE2D3 or mutants N77S, D87A (defective in closed conformation/thioester activation), and D117A (defective in lysine transfer). The RBR E3 retains activity with all mutants, supporting their utility as diagnostic tools to distinguish E3 ligases that require a closed E2~Ub conformations and those that facilitate allosteric ubiquitin transfer to lysine residues. **B)** Activity of $ZNFX1_{FL}$ WT and C1860A when partnered with an extended panel of UBE2D3 variants. E2 residues N77, D87, and L104 (UBE2D3 numbering) are essential for allosteric activation, as they promote and stabilize the closed E2~Ub conformation. In line with the necessity of a closed E2~Ub conformation for ZNFX1's E3 activity, substitutions at these key residues abolished activity, as evidenced by the absence of detectable signal. E3s requiring a closed E2~Ub conformation demonstrate a partial requirement for residue N114. $ZNFX1_{FL}$ demonstrates the same characteristic as a partial reduction in activity was observed. ZNFX1 is inactive with the E2 UBE2L3. This characteristic - an anomaly for a transthiolating E3 - is shared by the RING-Cys-Relay (RCR) E3 MYCBP2, where it can be accounted for by residue K96 (homologous residue is S94 in UBE2D3) undergoing steric clash with the MYCBP2 RING domain[2]. In support of the molecular basis for UBE2L3 incompatibility being related, $ZNFX1_{FL}$ activity was diminished when partnered with a UBE2D3 S94K variant. Residue S94 is otherwise functionally benign in UBE2D3, as the S94A variant exhibited activity comparable to the wild-type enzyme. **C)** Consistent with His1881 functioning as a general base that is essential for RZ transthiolating activity, it was ablated when $ZNFX1_{FL}$ H1881N was combined with UBE2D3 D117A. Activity could not be restored by increasing the basicity of the reaction buffer (pH 9.0). This suggests the $pK_a$ values for the ubiquitinated lysine residues are > 11 – underscoring the need for a catalytic base.

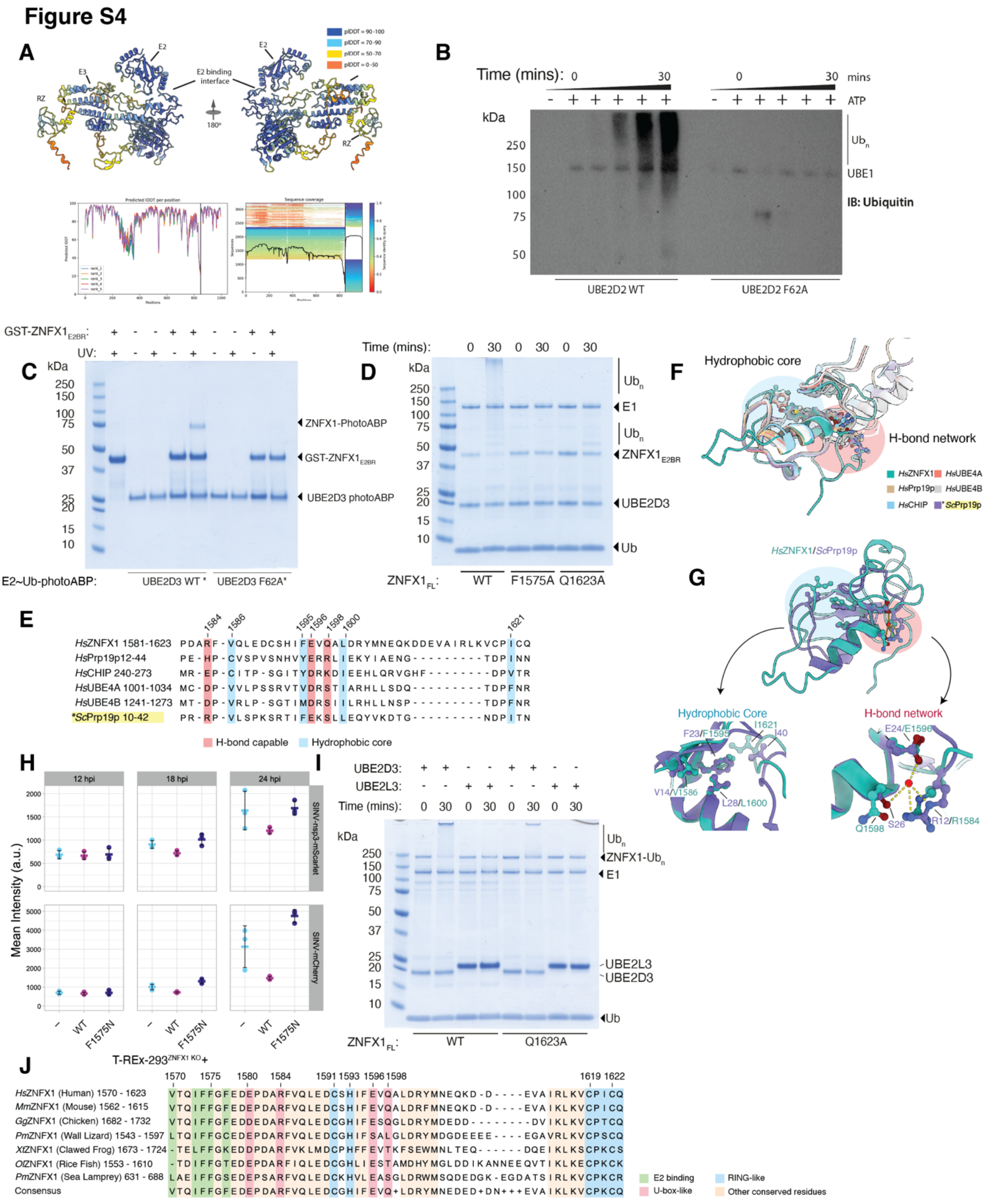

### Supplementary Figure 4

**A)** AlphaFold2 multimer model of ZNFX1 in complex with UBE2D3. The predicted complex is shown with both molecules in cartoon representation and coloured by pLDDT. The model highlights the putative E2-binding interface on ZNFX1. Structural predictions were generated using ColabFold with default settings. pLDDT scores for all models generated and sequence coverage is shown. **B)** ZNFX1$_{FL}$ autoubiquitination was assessed with WT and F62A UBE2D2.

Activity was observed only with WT, confirming that F62 is essential for functional interaction between the two enzymes. Autoubiquitination was detected by western blot using an anti-ubiquitin antibody. **C)** GST-ZNFX1$_{E2BR}$ (10 µM) underwent labelling with a UBE2D3 photocrosslinking UBE2D3~Ub probe (20 µM) selective for active allosteric E3 ligases [3]. Labelling was UV (365 nm LED lamp) and Phe62 dependent, further confirming the importance of this residue for functional interaction between ZNFX1 and UBE2D3. **D)** GST-ZNFX1$_{E2BR}$ variants (WT, F1575A and Q1623A) (1 µM) were tested for autoubiquitination. F1575A was inactive whilst Q1623A retained minimal activity. **E)** Structure-guided sequence alignment of the ZNFX1 Z-RING with a panel of human U-box E3 ligases and the yeast E3 *Sc*Prp19p reveals conservation at positions associated with the hydrophobic core and hydrogen bond (H-bond) network characteristic of U-box domains. The alignment was guided by structural similarity and secondary structure predictions using PROMALS3D. **F)** Structural alignment of the ZNFX1 Z-RING with representative U-box domains from the alignment in A). Residues structurally aligned with the U-box hydrophobic core and H-bond network are shown, along with the approximate location of these regions within the fold. **G)** ZNFX1 shares highest sequence similarity with the yeast U-box E3 ScPrp19p rather than human orthologues. In its Z-RING, Val1586, Phe1595, Leu1600, and Ile1621 align with hydrophobic core residues, while Arg1584 and Glu1596 match H-bonding residues. Although Ser26 of ScPrp19p is not conserved, ZNFX1 has Gln1598 at the equivalent position, potentially supporting hydrogen bonding. These features suggest that the Z-RING cross-brace may use water-mediated stabilization, as in U-boxes, instead of a second zinc ion typical of canonical RING domains. **H)** Red fluorescent signal in T-REx-293$^{ZNFX1\ KO}$ cells (blue) reconstituted with WT-(pink) or F1575N (purple) ZNFX1, and infected with SINV-nsp3-mScarlet (top panel) or SINV-mCherry (bottom panel) (MOI = 0.1). Fluorescence was measured at intervals of 15 min in an incubated plate reader (37°C and 5% $CO_2$). Data are presented as mean values ± SD from 3 independent infections. **I)** Autoubiquitination of ZNFX1$_{FL}$ WT and ZNFX1$_{FL}$ Q1623A was assessed with UBE2D3 and UBE2L3. Both variants were inactive with UBE2L3, as expected. With UBE2D3, Q1623A displayed substantially reduced activity relative to WT, suggesting Q1623 may act as a linchpin residue critical for optimal ligase function. **J)** Sequence alignment of the E2 binding region of ZNFX1 against a panel of vertebrates shows this region is highly conserved in ancestral ZNFX1 orthologs.

Figure S5

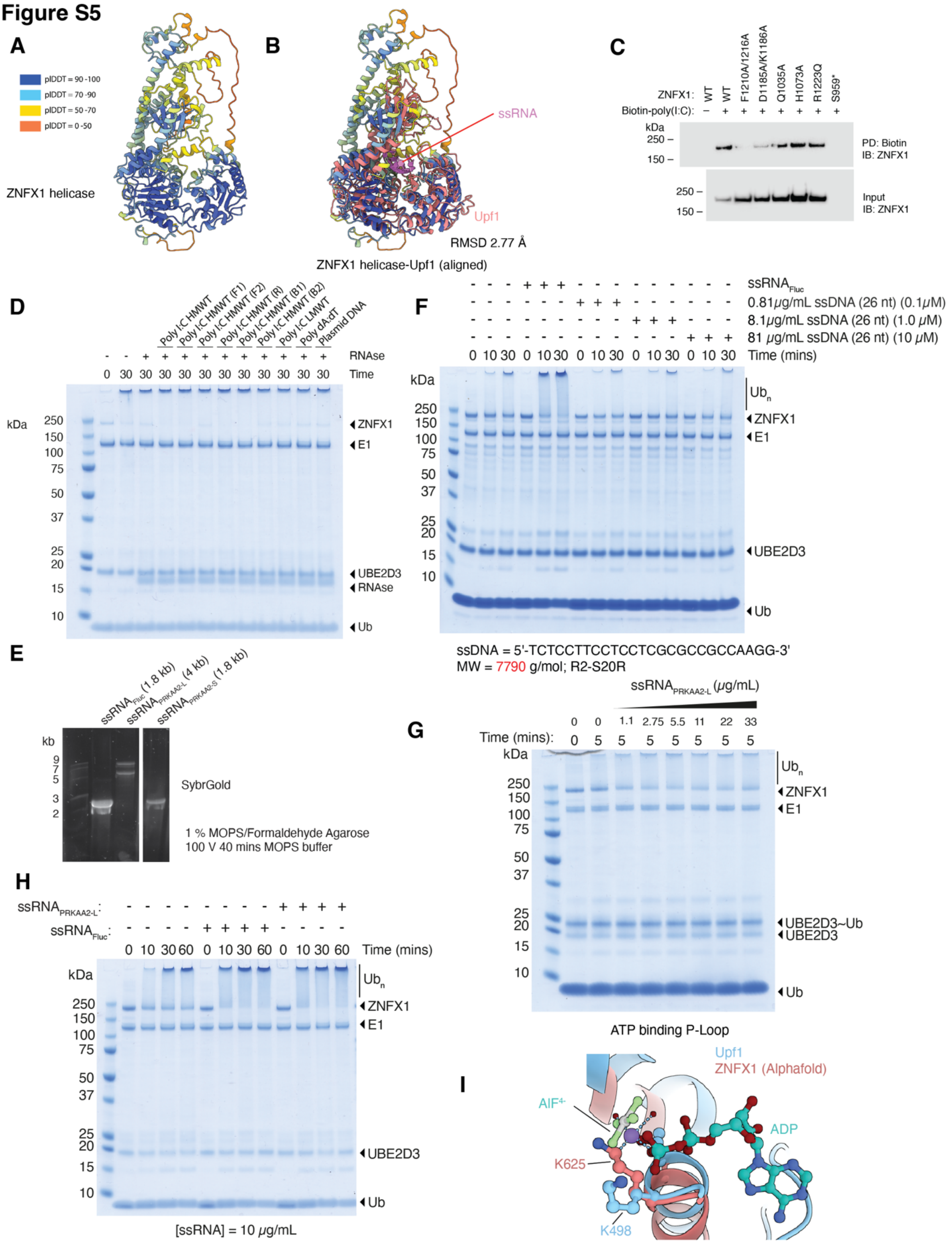


### Supplementary Figure 5

**A)** AlphaFold3 multimer model of the helicase region from ZNFX1 (residues 361-1259) full-length ZNFX1 Structural alignment of the ZNFX1 helicase domain with the Upf1–ssRNA complex. **B)** Experimental structure of human Upf1 bound to ssRNA (PDB: 2XZL) was aligned to the ZNFX1 model, revealing high structural conservation with an RMSD of 2.77 Å. ZNFX1 is coloured by pLDDTscore; Upf1 is shown in coral. **C)** Enrichment of WT or variant ZNFX1 by biotin-poly(I:C) from cell extracts. Western blots compared ZNFX1 abundance in input and enriched material. **D)** Nucleic acid-dependent activation of $ZNFX1_{FL}$. ZNFX1 was incubated with a panel of nucleic acids at 10 μg/mL: dsRNA species include high molecular weight poly I:C (HMWT), low molecular weight poly I:C (LMWT), biotin-, fluorescein- or rhodamine-labelled HMWT poly I:C (B1, B2; F1, F2; R, respectively). dsDNA species include double-stranded DNA (poly dA:dT), and circular plasmid DNA. Only the HMWT Poly I:C species (dsRNA) had an activating effect, implying ZNFX1 is selectively activated with large RNA molecules. **E)** Denaturing RNA gel for quality control of IVT products. ssRNA species were generated by IVT to test for ZNFX1 activation. An agarose (1 %), MOPS (10 %), formaldehyde (37 %) denaturing gel was used to evaluate the IVT products. Samples were heated at 70 °C for 15 min in formaldehyde loading buffer and stained with SYBR Gold. Although 1.8 kb $ssRNA_{Fluc}$ and 1.8 kb $ssRNA_{\textit{PRKAA2-S}}$ had reduced electrophoretic mobility compared to the 2 kb standard, they migrated equivalently, suggestive of a systematic discrepancy between the IVT products and the standard. The 4 kb $ssRNA_{\textit{PRKAA2-L}}$ ran as two distinct bands. This likely reflects incomplete denaturation and/or stable secondary structure formation. **F)** $ZNFX1_{FL}$ was tested with 1.8 kb $ssRNA_{Fluc}$ (10 μg/mL) or increasing concentrations of ssDNA (26 nt, 0.1, 1, and 10 μM). Activation was observed only with ssRNA, implying ssDNA of this size could not activate ZNFX1. **G)** Dose response of $ZNFX1_{FL}$ to ssRNA. Activation was most efficient between 2.75–11 μg/mL (2 – 8 nM) $ssRNA_{Fluc}$. A reduction in activity was observed at higher concentrations (22-33 μg/mL), suggesting a modest hook effect. **H**) ZNFX1 was incubated with either $ssRNA_{Fluc}$ or $ssRNA_{\textit{PRKAA2}\text{-L}}$ ssRNA species. Both induced comparable activation, indicating no significant size preference within this range. **I)** Conservation of the ATP-binding pocket in ZNFX1 and Upf1. Structural alignment of the AlphaFold3 ZNFX1 model with the Upf1–ssRNA complex (PDB: 2XZL) reveals conserved architecture of the ATP-binding site. The conserved lysine residue essential for ATPase activity in Upf1 is indicated (Lys498 in Upf1; Lys625 in ZNFX1). ZNFX1 is shown in coral, Upf1 in blue, and $ADP{\cdot}AlF_4^-$ is depicted in cyan and green/grey.

### Figure S6

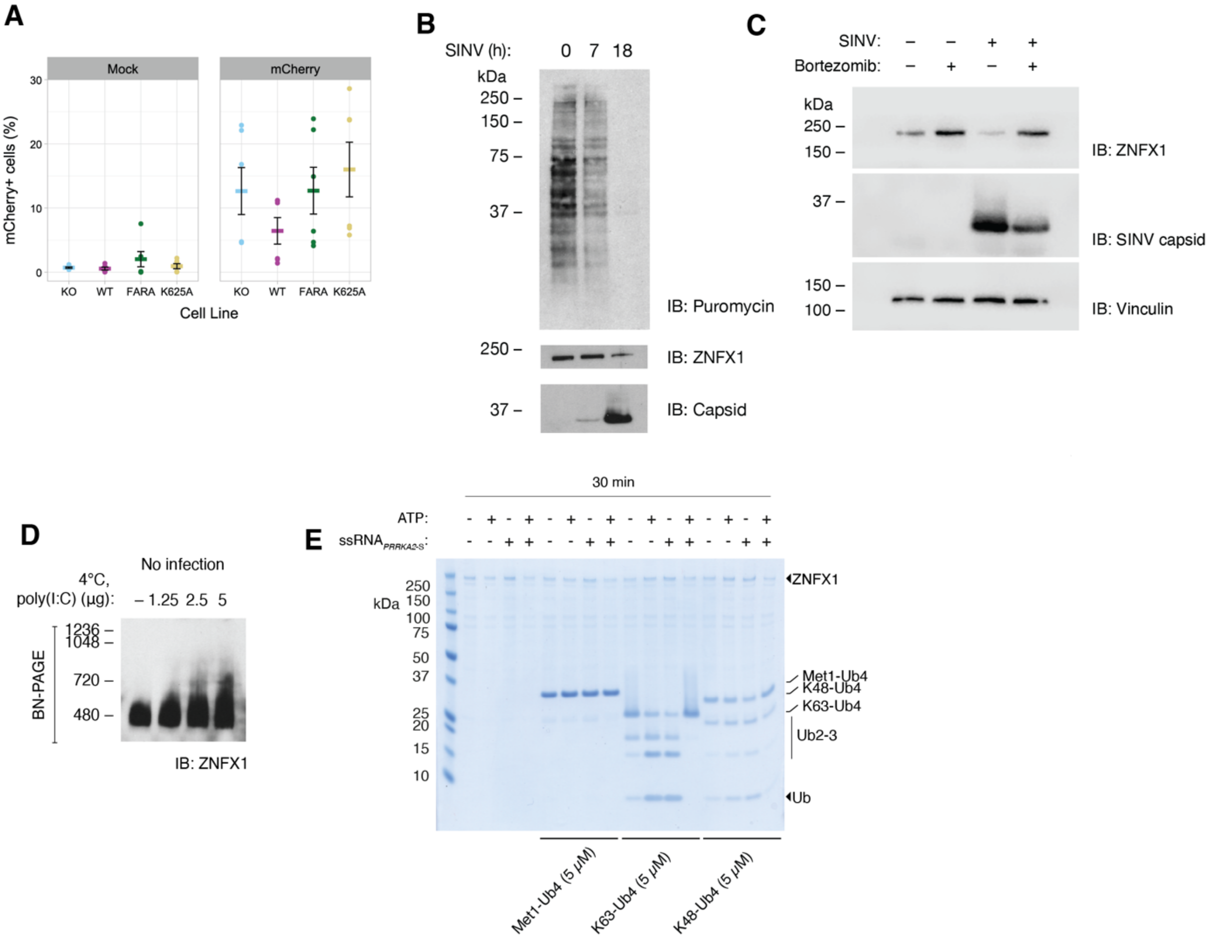


#### Supplementary Figure 6

**A)** Percent of cells mCherry+ cells following mock infection or infection with SINV-mCherry at an MOI of $3.75x10^{-3}$ for 24 h. Expression of FARA-ZNFX1 or K625A-ZNFX1 does not reduce the percent of SINV-infected cells. **B)** T-REx-293$^{ZNFX1\ KO}$ cells reconstituted with WT ZNFX1 were infected with SINV and treated with puromycin (5 ng/mL), which is incorporated into nascent proteins. Protein translation is halted at 18 h post-infection, as determined by protein puromycinylation visualised in Western blots. Probing for SINV capsid demonstrates the translation shut-off, and ZNFX1 disappearance, coincides with peak viral replication. **C)** T-REx-293$^{ZNFX1\ KO}$ cells reconstituted with WT ZNFX1 were infected with SINV (MOI = 1) and treated with the proteasome inhibitor Bortezomib (100 nM) for 18 h. Bortezomib treatment reduces ZNFX1 degradation and viral capsid protein, following infection. **D)** Blue Native PAGE Western blot of lysates from T-REx-293$^{ZNFX1\ KO}$ cells reconstituted with WT ZNFX1 treated with increasing doses of poly(I:C) for 10 min at 4 °C. **E)** Reaction presented in Figure 6F was carried out in the absence of E2~Ub. A methionine-linked (linear) ubiquitin tetramer was also included.

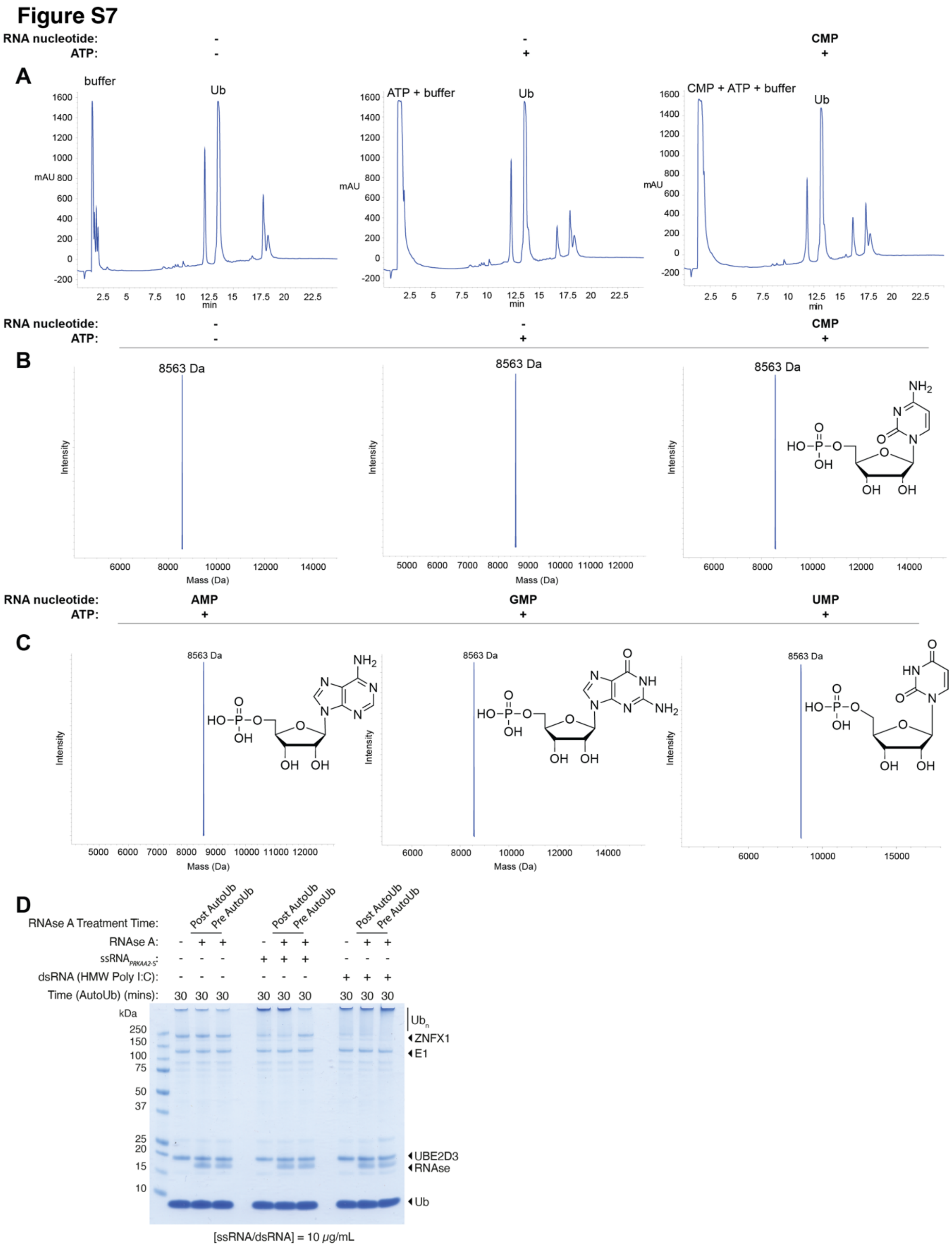


### Supplementary Figure 7

**A)** LC-MS UV chromatograms (214 nm) of representative ZNFX1 conjugation reactions containing E1 (0.4 μM), UBE2D3 (4 M), Ub (40 μM) ZNFX1 (1μM) and either no nucleotide and no ATP (left); no nucleotide and ATP (middle), and representative reaction for the addition of ATP and CMP (38 mM) (right). **B)** Deconvoluted mass spectra corresponding to reactions in E), showing a single species at 8,563 Da consistent with unconjugated ubiquitin (expected mass: 8,564.84 Da). **C)** Deconvoluted mass spectra of +ATP reactions with AMP (38 mM) (left), GMP (38 mM) (middle), and UMP (38 mM) (right), each also showing only the ubiquitin peak at 8,563 Da. No additional mass species corresponding to nucleotide-conjugated ubiquitin were observed under any condition. Reactions were incubated at 37 °C for 60 minutes and resolved with a 10-70% gradient (mobile phase A: $H_2O$ + 0.05% TFA; mobile phase B: acetonitrile + 0.04% TFA). **D)** ZNFX1 autoubiquitination in the presence of $ssRNA_{PRKAA\text{-}S}$ or dsRNA (poly(I:C)) is accompanied by RNA-dependent protein aggregation. RNase A treatment (100 µg/mL, 15 min, 37 °C) prior to autoubiquitination (30 min, 37 °C) prevented aggregation with $ssRNA_{PRKAA\text{-}S}$ but not with poly(I:C), consistent with RNase A resistance of the poly(I) strand. RNase A treatment after autoubiquitination had no effect on aggregate stability, indicating that RNA is required to initiate but not maintain ZNFX1 aggregation. Ubiquitin conjugation activity was inhibited prior to post-treatment RNase A addition using an E1-specific inhibitor.


1. Kamel, W., Ruscica, V., Embarc-Buh, A., de Laurent, Z.R., Garcia-Moreno, M., Demyanenko, Y., Orton, R.J., Noerenberg, M., Madhusudhan, M., Iselin, L., et al. (2024). Alphavirus infection triggers selective cytoplasmic translocation of nuclear RBPs with moonlighting antiviral roles. Mol Cell *84*, 4896-4911 e4897. 10.1016/j.molcel.2024.11.015.
2. Mabbitt, P.D., Loreto, A., Dery, M.A., Fletcher, A.J., Stanley, M., Pao, K.C., Wood, N.T., Coleman, M.P., and Virdee, S. (2020). Structural basis for RING-Cys-Relay E3 ligase activity and its role in axon integrity. Nat Chem Biol *16*, 1227-1236. 10.1038/s41589-020-0598-6.
3. Mathur, S., Fletcher, A.J., Branigan, E., Hay, R.T., and Virdee, S. (2020). Photocrosslinking Activity-Based Probes for Ubiquitin RING E3 Ligases. Cell Chem Biol *27*, 74-82 e76. 10.1016/j.chembiol.2019.11.013.
